# Retrotransposon expression is upregulated by aging and suppressed during regeneration of the limb in the axolotl (*Ambystoma mexicanum*)

**DOI:** 10.1101/2024.04.04.588033

**Authors:** Samuel Ruiz-Pérez, Karla Torres-Arciga, Cynthia Gabriela Sámano-Salazar, José Antonio Ocampo-Cervantes, Alejandra Cervera, Clementina Castro-Hernández, Ernesto Soto-Reyes, Nicolas Alcaraz, Rodrigo González-Barrios

## Abstract

The axolotl (*Ambystoma mexicanum*) has a great capacity to regenerate its tissues and whole-body parts; however, the fidelity and success of its regenerative process diminish with age. Retrotransposons make up the largest portion of the axolotl genome, and their expression may be involved in this age-related decline. Through an integrative analysis of repetitive element expression using RNA-seq, we show that Ty3 retrotransposons are highly upregulated in the axolotl as an effect of chronological aging. Other non-LTR transposons, including LINE-1, function as hubs of gene coexpression networks involved in muscle development, N-methyltransferase activity, amyloid proteolysis, and regulation of apoptosis and connective tissue replacement, which are also suppressed by the increase in age. In contrast, we find that during regeneration of the limb these pathways and the expression of Ty3 retrotransposons are distinctly downregulated. Although the blastema is able to readjust most of the transposon dysregulation caused by aging, there are still several elements that remain affected and may have an impact in the metabolic and immune responses during the regenerative process. We also report that numerous C2H2-ZFPs, especially KRAB-ZPFs, are coexpressed with hub retrotransposons, which reveals their potential as important regulators of transposable element expression during aging and regeneration. This integrative analysis provides a comprehensive profile of retrotransposon expression through chronological aging and during limb regeneration in the axolotl and indicates that transposable elements are responsive to physiological changes in a tissue-specific way and participate in the gene co-regulatory networks underlying the regenerative process.

**Graphical abstract.:** Transcriptomic analyses of adult and sub-adult axolotls offer an insight into the potential role of retrotransposons during limb regeneration and how they are affected by chronological aging. Retrotransposons of the Ty3 superfamily are usually suppressed during regeneration, but after experiencing a significant upregulation through the aging of the axolotl, their regulation during tissue repair is limited. It has been reported that the success of the axolotl’s regenerative process diminishes with age. Our findings show that Ty3 and other transposon families, such as LINE-1, are involved in the gene regulatory networks that suppress muscle development, apoptosis control, and tissue replacement due to aging. These results suggest that repetitive element expression may indirectly constrain the regenerative capacities of older axolotls.

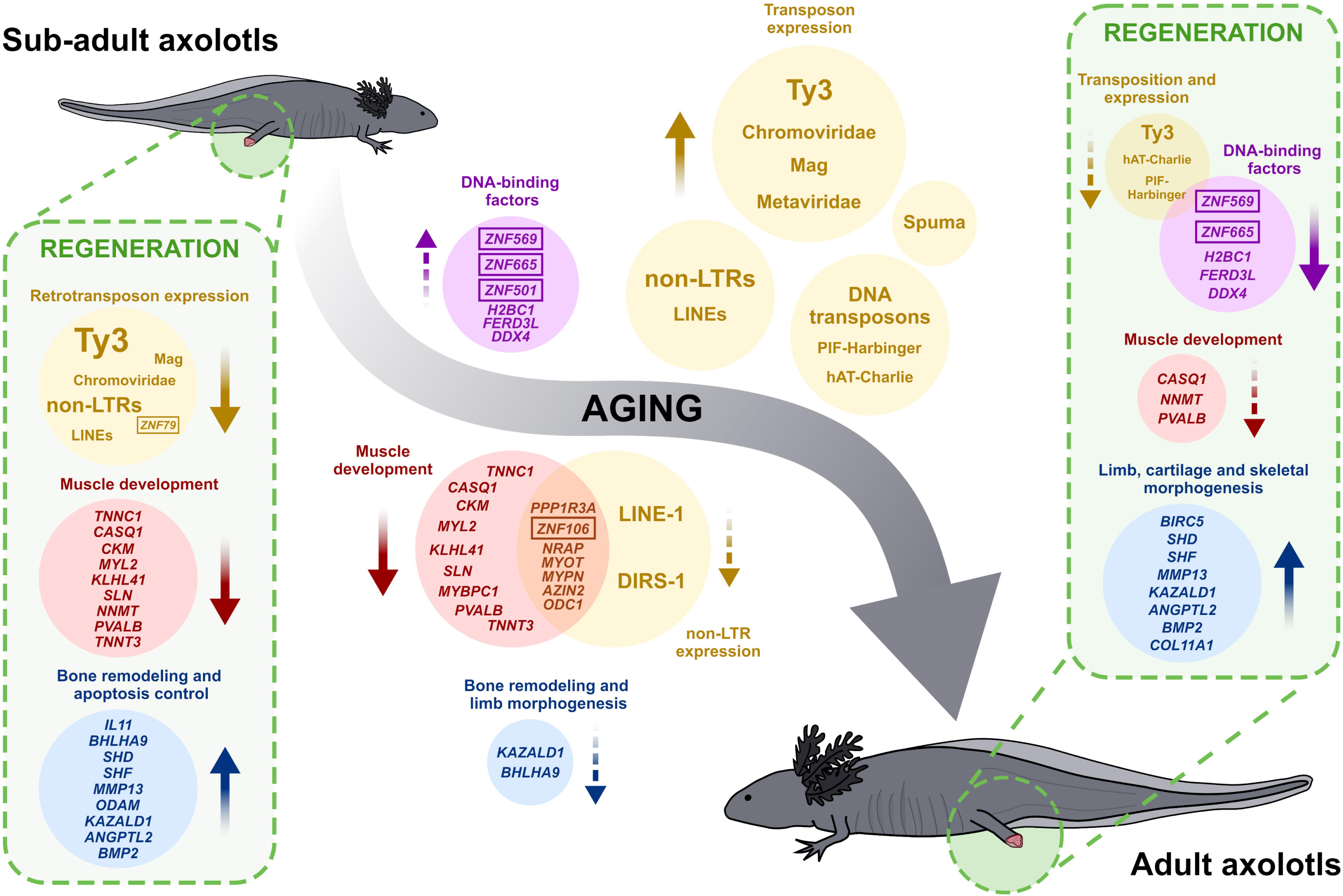

## Introduction

The Mexican axolotl or *Ambystoma mexicanum* (Shaw & Nodder, 1789) [1] is a salamander native to the canals and lakes of Xochimilco and Chalco, Mexico, that has been widely studied due to its great capacity to regenerate its cells, tissues, and whole-body parts, including limbs, heart, brain, and lungs.

Although *A. mexicanum* retains a remarkable regenerative capacity throughout its life that far exceeds that of humans and other mammals, the rate, fidelity, and success of its regenerative process diminish with age [2]. Among the many factors that could influence this decline are the increase in body and extremity size and mass, ossification of the skeleton, thickening and loss of flexibility of the skin, changes in nerve function, and shifts in circulating factors, hormones, immune response, or metabolism [2]. However, the causes of this age-related decrease in its regenerative capacity have not been fully clarified.

Until recently, research into the factors underlying regenerative biology has been mostly characterized by studies of protein-coding genes [3]. However, in the case of the axolotl, the numerous repetitive elements (REs) within its ∼32 Gb genome are particularly interesting as molecular interactors with a potential role during regeneration. Based on the most recent genome sequence assembly for the d/d strain (AmexG_v6.0-DD), around 65 % (∼18.6 Gb) of the axolotl genome consists of repetitive sequences [4]. Moreover, the largest portion of the axolotl repeatome is composed of long terminal repeat (LTR) retroelements (∼26 %, ∼7.2 Gb) of the Ty3 type (∼11%, 3.2 Gb).

Several studies have reported the negative effects of repetitive element activity at both the genomic and physiological levels. For example, retrotransposition can introduce potentially catastrophic alternative gene splicing patterns and gene expression profile changes in some diseases [5]. In such cases, silencing of active REs could be a critical part of the regulation and maintenance of essential cellular homeostasis. In contrast, other studies have shown that the activity of some REs, especially transposable elements (TEs), can be beneficial to their hosts [3,6]. For example, by regulating the expression of protein-coding genes at the transcriptional and post-transcriptional levels, retrotransposons can perform important functions in embryogenesis, such as affecting pluripotency and directing cell fate decisions [3,7]. Nonetheless, the involvement of REs in animal regeneration has been addressed by only a few studies in not many model systems [3,8–10].

For instance, Mashanov et al. [8] have reported large-scale changes in the transcriptional activity of LTRs during regeneration of radial organ complexes in the sea cucumber *Holothuria glaberrima*. Via high-throughput transcriptomic analyses, the authors identified 36 LTR retrotransposons belonging to Bel and Ty3 clades, among which 20 LTRs significantly changed their expression level during regeneration. The most significantly up-regulated element, Ty3-1_Hg [11], showed a drastic (>50-fold) increase in expression in glia and neurons of the regenerating central nervous system [3,8]. Moreover, the increases in retrotransposon transcription reportedly did not have a deleterious effect on genomic integrity nor resulted in programmed cell death, but, instead, contributed to regeneration.

In the case of the axolotl, Zhu *et al.* [9] reported both overexpression and higher retrotransposition of a long interspersed nuclear element 1 (LINE-1) in the dedifferentiating tissues of the limb blastema. Using qPCR, they showed that there was a dramatic upregulation of LINE-1 transcriptional activity as early as 2 days post-amputation (dpa). They also reported that other putative TEs and LINE-1-related piRNAs appear to be expressed or transcriptionally activated in the regenerative limb, with different dynamics. Even though the authors did not propose a precise function for LINE-1 or other repetitive elements in regeneration, they suggested that their reactivation is a ubiquitous phenomenon and may serve as a marker for cellular dedifferentiation in the early-stage of limb regeneration.

Although the studies in both *A. mexicanum* and *H. glaberrima* have demonstrated differential expression of retroelements during regeneration, their interpretations of the specific response and function of REs after a traumatic injury is still contrasting. One possibility is that the transcriptional activity of retrotransposons in the regenerate is a non-specific consequence of an overall reduction of epigenetic silencing, which in the case of the axolotl could allow the dedifferentiating cells of the blastema to revert to and establish a unique germline-like state [9]. In such case, transposons could also be making use of the global injury-induced chromatin activation and transcriptional de-repression caused by the dedifferentiation process to amplify themselves and thereby increase their odds of surviving the lysis of their host cell and being taken up by other cells [3,12]. In contrast, Mashanov *et al.* [3] suggest that retrotransposon expression changes during regeneration might not represent a mere exploitation of the transcriptional machinery of the host with aims of propagation, but rather that this dysregulation of transcription might be specifically controlled by the host at the tissue, cell, and repetitive element levels.

Taken together, these findings indicate that REs are specifically controlled by the host, respond in a coordinated way to regeneration cues and thus may play some previously unrecognized roles in post-traumatic tissue regrowth [3]. In this context, the physiological state of the organism could also be a crucial factor with an impact on the coordinated function of retrotransposons during regeneration. Two main questions have been proposed regarding retroelements’ role in the regenerative process: 1) what causes the change in their expression levels?, and 2) what are the repercussions of this dysregulation on the host organism? [3]. On top of that, we reasoned that, in the axolotl, the expression profiles of repetitive elements may be modified by a physiological or pathological process (such as aging) before an injury and outside of the regenerative process, which could then modify the basal state from which the organism responds to tissue damage and regulates regeneration. Repetitive element transcriptional dysregulation might be a cause or effect of the shifts in circulating factors, hormone production, immune response, or metabolism, that potentially lead to the age-related decrease in regenerative capacity in the axolotl.

In fact, studies in mice and other model organisms have shown that active transposable elements contribute to the aging process [13,14], and both active and inactive elements also accumulate in age-related neurodegenerative processes and diseases [15]. Also, RE de-repression has been described in senescent cells [16], and RE transcripts have been implicated in inflammation and oxidative stress [17], two key contributors to aging and disease with positive and negative effects for regeneration. For example, recent work shows that LINEs become active with aging and promote senescence-associated inflammation, making LINE-1 reverse transcriptase a relevant target for the treatment of age-associated disorders [13]. The underlying mechanisms by which RE transcripts affect these processes are not yet clear, but could involve activation of innate immune responses, which also play a central role in tissue repair and regeneration [18]. As such, changes in global RE transcript levels have been reported to be a better marker of biological age than protein-coding genes in both human fibroblasts and *Caenorhabditis elegans* [19]. One reason for this could be that RE expression changes occur consistently and in the same direction, whereas gene expression patterns may increase or decrease variably with age-related processes [19]. In summary, the role of repetitive elements amid the interplay of aging and reparative regeneration is still unclear. However, many studies so far have focused on known coding genes and overlooked repetitive or non-coding genetic material, which is coincidentally the majority of the genome in the case of the axolotl.

In this study, we conducted RNA sequencing analysis to characterize the transcriptomic profiles of protein-coding genes and repetitive elements in the regenerating limb of native (Xochimilco) wild-type Mexican axolotls from two different age groups. In addition, to discern if gene and RE expression is affected differently by age in distinct tissue types, we integrated into the analysis 124 RNA-seq samples from previously published studies comprising non-limb, limb, and limb blastema tissues divided into sub-adult and adult age groups. We report that repetitive elements, mainly of the Ty3 superfamily, are predominantly upregulated as an effect of chronological aging, and also downregulated during regeneration of the limb. The blastema of adult axolotls is able to counterregulate most but not all the retrotransposon expression affected by an increase in age. Additionally, we identified several coexpression modules that suggest an indirect TE-mediated regulatory network directing the immune, metabolic, and developmental responses during aging and regeneration.

## Results

### A major suppression of muscle development and function underlies limb regeneration

To determine the genes that are differentially expressed by aging and during limb blastema formation, we extracted RNA samples from the amputation of a posterior limb of five wild-type male sub-adult axolotls (aged 8 months) and two wild-type male adult axolotls (aged 8 years). Ten days after amputation, we collected blastema tissues only from the five sub-adult axolotls (**Fig 1A**), since none of the adult axolotls displayed any development of regenerative tissue even after 6 months of monitoring. After extraction, we subjected the samples to RNA-seq, and quantified and analyzed their gene and RE expression levels following two distinct pipelines (see **Materials and Methods** and **S1 Fig**).

**Fig 1.**
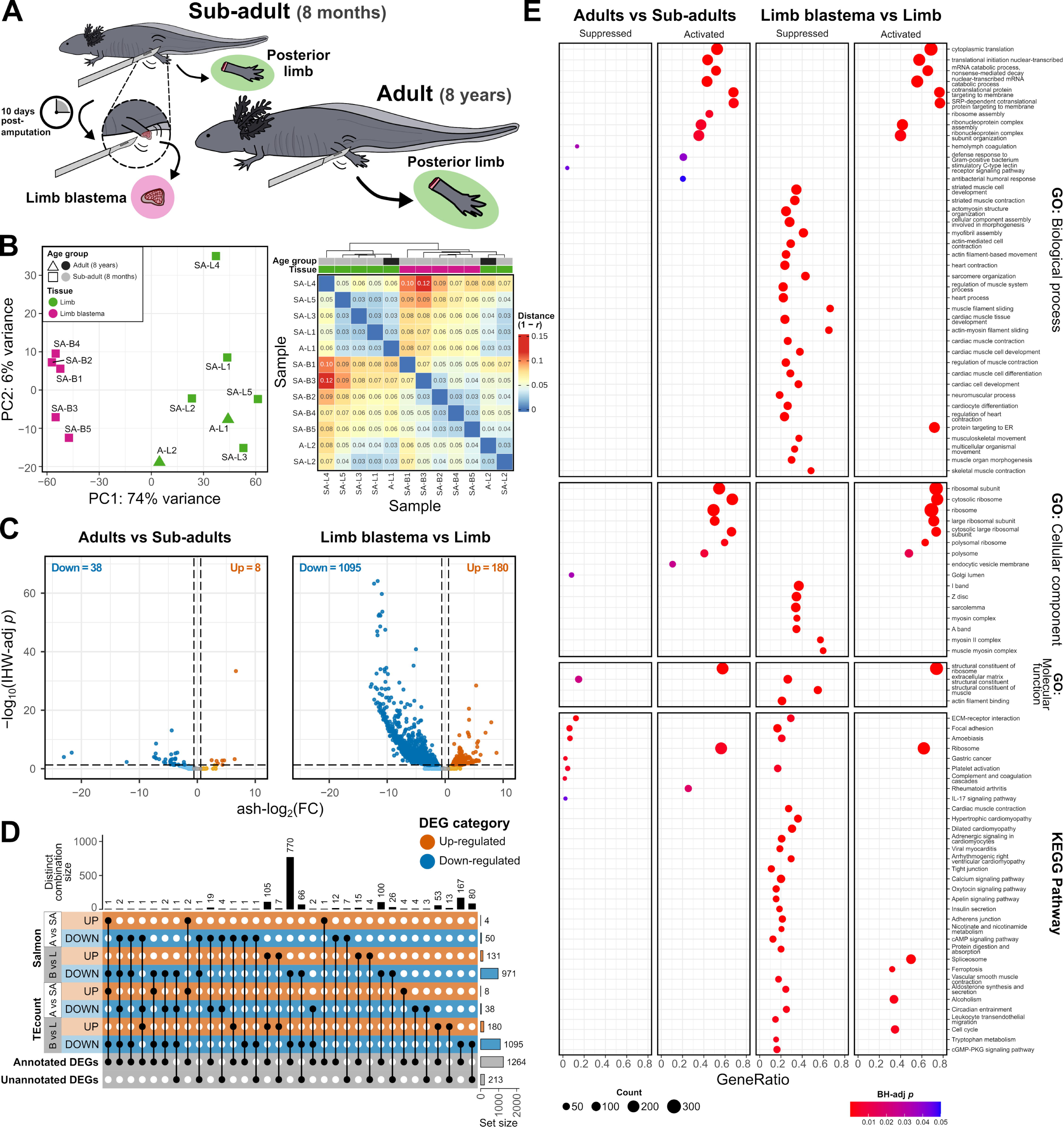
Gene expression differences between native axolotl samples of distinct age groups and tissues. (A) Graphical representation of the tissue sampling of native axolotls, which consisted of five limb amputations and five limb blastemas (10 days post-amputation) from sub-adult specimens (8 months old), and two limb amputations from adult specimens (8 years old). **(B)** Left: principal component analysis (PCA) plot of gene and repetitive element counts after variance stabilizing transformation (VST). Right: heatmap of sample-to-sample distances (1 − Pearson correlation(VST[counts])). **(C)** Volcano plots show the gene expression differences between adult and sub-adult samples and between blastema and limb samples. Genes with significantly increased (ash-log_2_(FC) > log_2_(1.5), IHW-adj *p* < 0.05) or decreased (ash-log_2_(FC) < −log_2_(1.5), IHW-adj *p* < 0.05) expression (DEGs) are represented by dark orange and dark blue points, respectively. **(D)** UpSet plot of the total DEGs per contrast and quantification method, their distinct combinations, and their annotation status. **e)** Dotplot shows the significantly activated or suppressed gene ontology (GO) terms and Kyoto Encyclopedia of Genes and Genomes (KEGG) pathways identified for each contrast by fast gene set enrichment analysis (FGSEA). SA: sub-adult, A: adult, L: limb, B: limb blastema.

Principal component analysis (PCA) of the normalized expression counts revealed the presence of two underlying subgroups within the dataset, corresponding to the two tissue types: limb and limb blastema (**Fig 1B**). Based on Pearson’s correlation, a sub-adult limb tissue (SA-L4) showed the highest sample-to-sample distance when contrasted with two sub-adult blastemas (SA-B3 and SA-B1) (**Fig 1B**).

Differential expression analysis (DEA) revealed that 971–1095 of the differentially expressed genes (DEGs) in the contrast between limb blastema and limb tissue were downregulated, and 131–180 genes were upregulated (**Fig 1C**). Importantly, in total for all contrasts, 213 genes are not annotated in the *A. mexicanum* genome reference assembly v6.0-DD (**Fig 1D**). Next, we conducted gene set enrichment analysis to identify which biological functions or processes were enriched among all annotated genes. In accordance with DEA results, the blastema vs limb contrast also showed a higher downregulation at the process level, with the suppression of 598 GO terms and 47 KEGG Pathways and the activation of 303 GO terms and 17 KEGG Pathways (**Fig 1E**). We found that the suppressed terms were mainly associated with muscle cell development and differentiation, muscle contraction, muscle component assembly and organization, muscle signaling pathways, extracellular matrix components and signaling, and cellular junctions. In contrast, activated processes were primarily related to ribosomal components and processes, translation, mRNA catabolism and protein targeting.

### Native adult axolotls display an activation of RNA and ribosomal processes

To identify the genes and biological processes generally impacted by chronological aging (an 88-month age difference), we contrasted the tissue samples from adult axolotls against the samples from sub-adult axolotls. Even though the total number of DEGs in this contrast was lower than the blastema vs limb comparison, it similarly showed more downregulated than upregulated genes (**Fig 1C**). Conversely, most enriched GO terms were activated rather than suppressed, albeit 7 KEGG Pathways were suppressed and 2 were activated. The top suppressed terms were related to extracellular matrix constituents, coagulation, the Golgi lumen, and the stimulatory C-type lectin receptor signaling pathway. As in the blastema vs limb contrast, activated terms in the age contrast included ribosomal element assembly, translation, mRNA catabolism and protein targeting.

### Ty3 retrotransposon subfamilies expression is downregulated during limb regeneration

Next, we quantified and analyzed RE expression at the subfamily level. Repetitive element “insertions” or loci within a particular genome can be grouped into subfamilies, which are sets of REs that are highly related at the sequence level and are relatively distinct from other elements [20]. In the comparison of the limb blastema against the limb tissue, DEA revealed only 7–8 significantly differentially expressed RE subfamilies (DEREs), i.e., 2–6 downregulated and 1–6 upregulated, although there was an apparent pattern of predominant RE downregulation (**Fig 2A and 2B**). Among these DEREs, 6 belong to the Chromoviridae family and 6 to the Mag family, both members of the Ty3 retrotransposon superfamily and the LTR class. Three DEREs have not been annotated in the reference RE library for *A. mexicanum* and are labeled as pertaining to an “Unknown” class. For the age group contrast only one quantification method, ExplorATE, allowed the detection of DEREs (**Fig 2B**).

**Fig 2.**
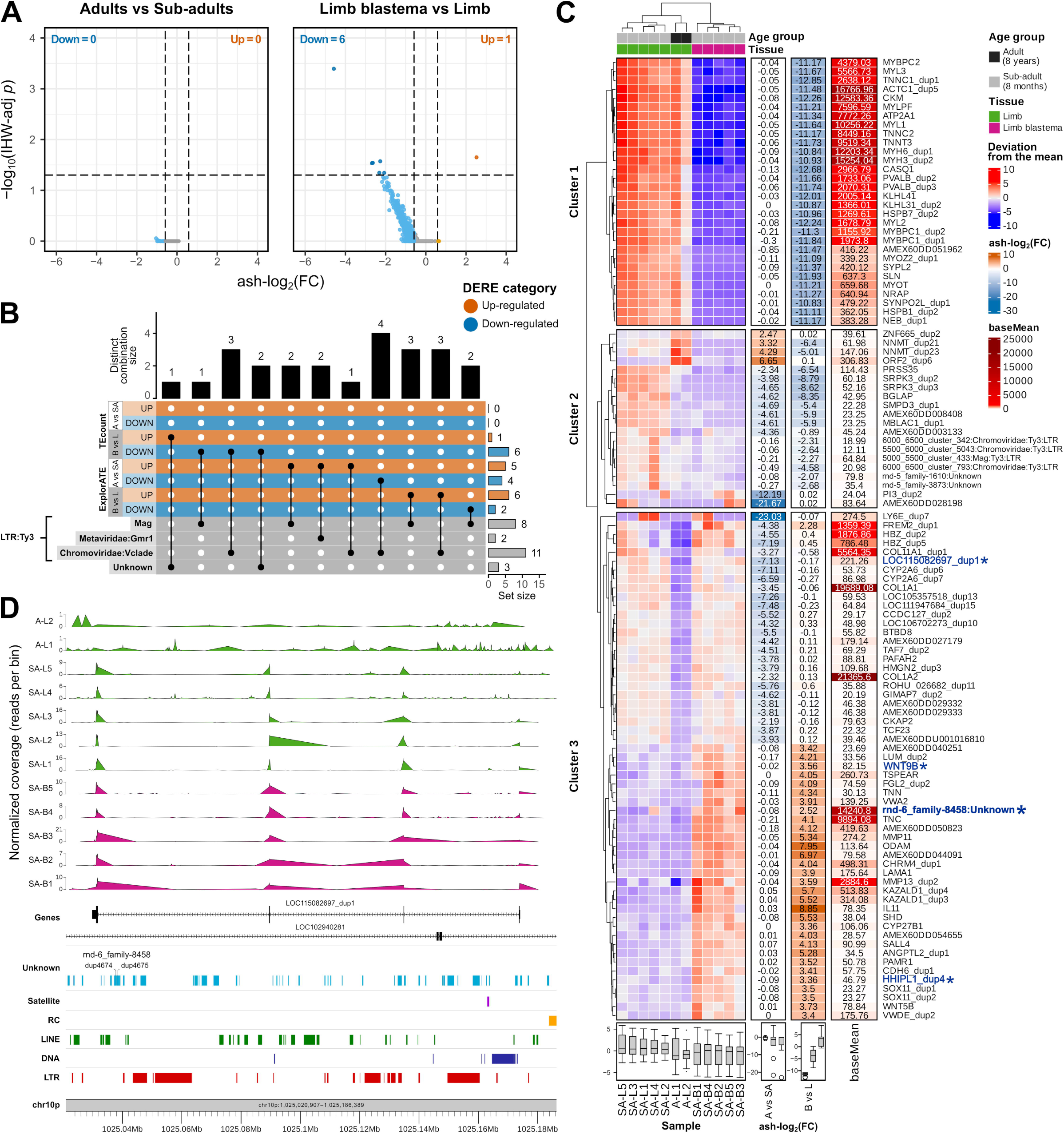
Repetitive element expression changes in the axolotl’s regenerating limb. (A) Volcano plots show the repetitive element (RE) expression differences between adult and sub-adult samples and between blastema and limb samples. RE with significantly increased (ash-log_2_(FC) > log_2_(1.5), IHW-adj *p* < 0.05) or decreased (ash-log_2_(FC) < −log_2_(1.5), IHW-adj *p* < 0.05) expression (DEREs) are represented by dark orange and dark blue points, respectively. **(B)** UpSet plot of the total DEREs per contrast and quantification method, their distinct combinations, and their annotation status. **(C)** Heatmap of the deviation from the mean of each VST-normalized gene/RE count, split into clusters by the global dendrogram. Columns with ash-log_2_(FC) by contrast and baseMean are shown on the right for each gene/RE. Genes labeled in blue and with an asterisk (*) have a genomic overlap or relative proximity to a locus of the RE, also marked in blue with an asterisk. **(D)** Karyoplot of *LOC115082697_dup1*, one of the three differentially expressed genes in the blastema that overlap with loci of a DERE in the same heatmap cluster. RNA-seq read coverage of each sample is plotted on top. SA: sub-adult, A: adult, L: limb, B: limb blastema.

To jointly compare the expression of genes and REs, we performed hierarchical clustering on the elements with the highest absolute log fold change (LFC) for both the tissue and the age group contrasts. To this end, we calculated the amount by which each gene or RE deviated from its mean across all samples and then split the selected genes and REs into three clusters by the global dendrogram (**Fig 2C**).

We found two main blocks of genes/REs that covary across tissue types (blastema vs limb), one includes downregulated genes related to muscle development and function (cluster 1: *MYL2*, *MYH3*, *CASQ1*, *TNNT3*, etc.) and the other connects an unknown RE (*rnd-6_family-8458*) with upregulated genes related to bone formation, remodeling and regeneration (bottom cluster 3: *KAZALD1*, *ODAM*, *IL11*, *ANGPTL2*, etc.). In regard to the age group contrast, we found another set of genes for which the adult axolotls have both lower expression (top cluster 3: *LY6E*, *HBZ*, *COL1A1/2*, *LOC115082697*, etc.) and higher expression (top cluster 2: *ORF2*, *NNMT* and *ZNF665*) than the sub-adult axolotls. Conversely, we found a set of genes/REs that covary across both tissues and age groups: a serine/arginine-rich protein-specific kinase (*SRPK3*), an inactive serine protease (*PRSS35*), osteocalcin (*BGLAP*), a sphingomyelin phosphodiesterase (*SMPD3*) and an endoribonuclease of histone-coding pre-mRNA 3’-end (*MBLAC1*). These genes, together with four Ty3 retrotransposons and two unknown REs, are also downregulated in blastemas when compared to limb tissues.

Repetitive elements have shown considerable regulatory potential, specifically in the modification of gene expression. To address the possible role of the DEREs identified in the blastema on gene regulation, we evaluated their genomic context and vicinity to DEGs. To do this, we annotated each locus of each DERE to its nearest genomic/genic feature. As shown in **Table 1**, most DERE loci were located in intergenic regions and in the introns of non-DEGs. Furthermore, we found that only one unknown DERE (*rnd-6_family-8458*), contained loci that both colocalized (had a genomic overlap or relative proximity) and shared a deviation-from-the-mean cluster with a DEG (i.e., *WNT9B*, *HHIPL1*, *LOC115082697*). Specifically, only 2 out of 5724 loci of *rnd-6_family-8458* are contained within the most downstream intron of *LOC115082697*, a gene that is underexpressed in adult axolotls (**Fig 2C and 2D**). Interestingly, our searches of the translated consensus sequence of *rnd-6_family-8458* against the InterPro 98.0, PFAM 36.0 and RepeatPeps (RepeatMasker) databases only revealed one match: a region of a protein which is predicted to be embedded in the membrane (TMhelix). These results suggest that the potential regulation of gene expression by REs in the blastemas of native axolotls is independent of their genomic proximity to the genes.

**Table 1.**
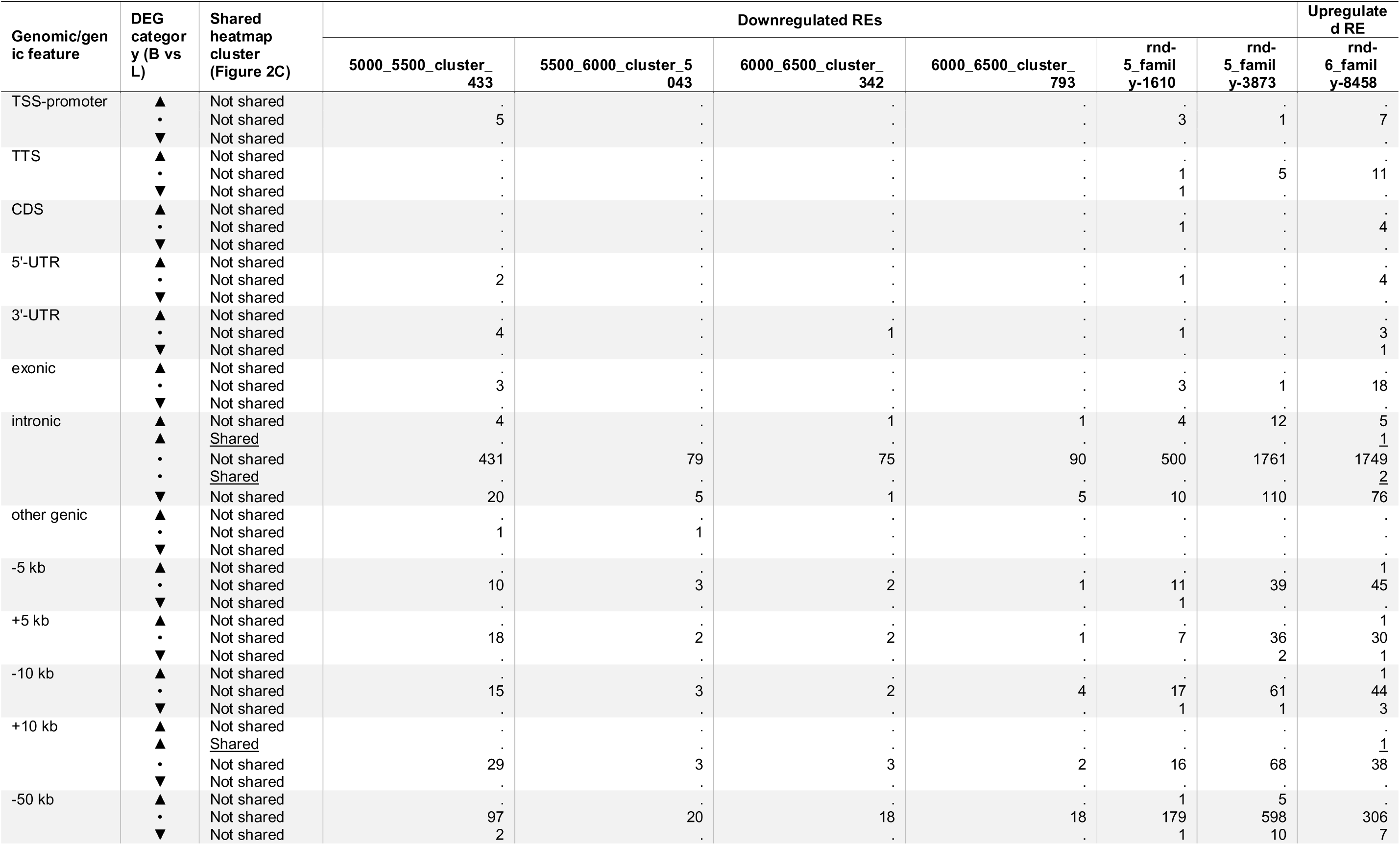
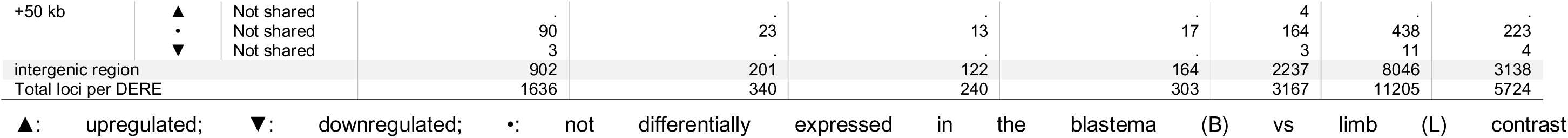
Summary of the loci of differentially expressed repetitive element subfamilies and the genomic features (gene subfeatures or intergenic regions) with which they overlap for the blastema vs limb contrast of axolotl tissues.

### Age-related muscle development downregulation in the limb replaces muscle suppression in the blastema

To determine if the expression of genes with roles in regeneration is affected differently by age in distinct tissue types, such as the blastema compared to the limb, we integrated 124 previously published RNA-seq samples (**S1 Table**) [4,21–23] that included non-limb, limb, and limb blastema tissues, divided into the sub-adult and adult age groups. We then quantified and contrasted their gene and RE expression levels together with the samples of this study. Based on the PCA of all 136 samples (**Fig 3A**), we conducted the subsequent DEAs considering batch effects.

**Fig 3.**
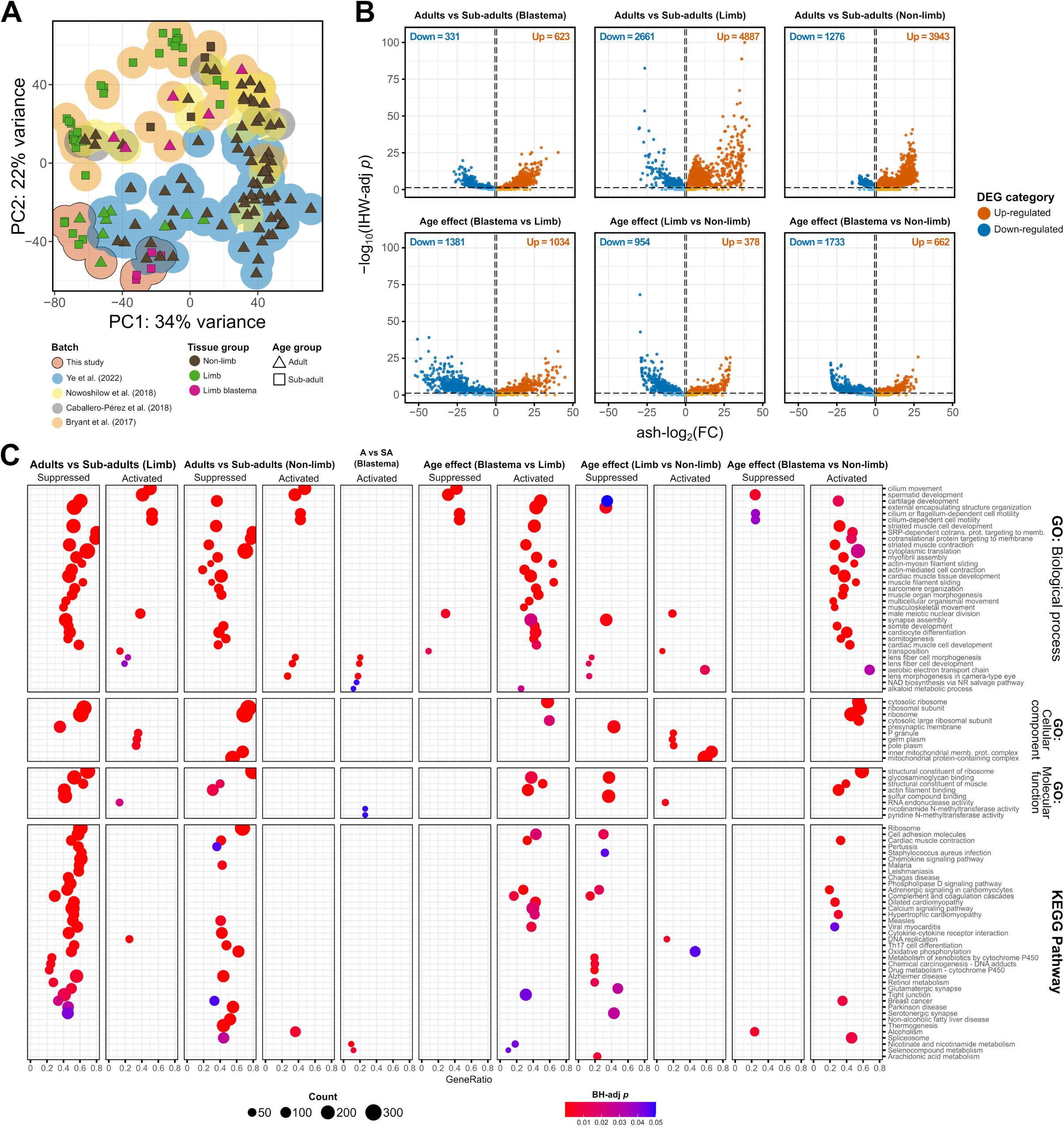
Gene expression differences between global axolotl samples of distinct age groups. (A) Principal component analysis (PCA) plot of gene and repetitive element counts after variance stabilizing transformation (VST). **(B)** Volcano plots show the gene expression differences between adults and sub-adults for each tissue type and the age effect differences in each pairwise comparison of tissues. Genes with significantly increased (ash-log_2_(FC) > log_2_(1.5), IHW-adj *p* < 0.05) or decreased (ash-log_2_(FC) < −log_2_(1.5), IHW-adj *p* < 0.05) expression (DEGs) are represented by dark orange and dark blue points, respectively. **(C)** Dotplot shows the significantly activated or suppressed gene ontology (GO) terms and Kyoto Encyclopedia of Genes and Genomes (KEGG) pathways identified for each contrast by fast gene set enrichment analysis (FGSEA). SA: sub-adult, A: adult, L: limb, B: limb blastema.

In brief, we found that older axolotls exhibit many underexpressed and overexpressed genes compared to their younger counterparts in all tissues evaluated (**Fig 3B**). We identified 3943–4281 upregulated and 1190–1276 downregulated genes in the contrast of non-limb tissues, 4763–4887 upregulated and 2490–2661 downregulated genes in the comparison of limbs, and 623–748 upregulated and 331–421 downregulated genes in the blastemas contrast. Moreover, 2622 (∼54 %) of the 4887 genes that were upregulated in the contrast of adult vs sub-adult limbs were also downregulated in the adult blastema, which represents a ∼63 % of the 4176 total genes downregulated in adult blastemas when contrasted with adult limbs (**Figs 3B and S2A**). However, in terms of both quantity and type of enriched biological pathways, non-limb and limb tissues from adult axolotls are much more similar to each other than to the adult blastema: chronological aging in the limb and non-limb tissues triggers a substantial suppression of pathways related to translation and muscle development and the activation of cell motility and cilium or flagellum formation. Meanwhile, in the blastema age only prompts a limited activation of pathways related to lens fiber cell development and nicotinate, nicotinamide, pyridine, and seleno-compound metabolisms (**Fig 3C**).

In accordance with what we observed in the age contrasts, pairwise tissue contrasts revealed that during regeneration, the blastema exhibits both a major downregulation and a minor upregulation of genes compared to the limb and non-limb tissues (**S2A Fig**). At the biological pathway level, the blastemas from adult axolotls show significant activation of the cytosolic ribosome, embryonic limb and skeletal system morphogenesis, and non-embryonic limb and cartilage development, when compared to the limb (**S2B Fig**). At the same time, they exhibit suppression of transposition, cilium or flagellum-dependent cell motility and spermatid development. The underexpression and overexpression of genes between the blastema and other tissues are both stronger in samples of adult axolotls than in samples from sub-adult axolotls (**S2A Fig**). However, blastemas from sub-adult axolotls do not exhibit significant pathway activation in comparison with limb tissues, but only a suppression of cell components involved in morphogenesis and pathways of muscle development and contraction, such as myofibril assembly and sarcomere organization (**S2B Fig**).

In summary, we detected a significant difference in the effect of age on the blastema, compared to the effect it has on the limb. Namely, that relative to the limb, the increase in age has a net positive effect on striated muscle, cartilage, and synapse development in the blastema (**Fig 3C**). In other words, the age-related suppression of muscle and nervous structure development observed in the aged limb, appears to reach an almost equivalent effect, outside of the regenerative process, to the muscle suppression needed during the development of the blastema. In contrast, since the limbs from sub-adult axolotls have not undergone an age-related effect, their blastemas do induce the downregulation of striated muscle development (**S2B Fig**).

### Aging axolotls’ limbs undergo a Ty3 retrotransposon upregulation that is absent during regeneration

Subsequently, we evaluated the expression of REs at the subfamily level in all samples. We found that older axolotls exhibit a higher expression of retrotransposons than their younger counterparts in the extremity and other tissues, but not in the regenerative tissue of the limb. In the comparison of adult and sub-adult non-limb tissues, we identified 3–31 downregulated and 134–396 upregulated DEREs, with most being Ty3 retrotransposons (LTRs) or unknown REs (**Fig 4A and Table 2**). Similarly, the contrast of limb tissues of different age groups showed 0–24 downregulated and 223– 519 upregulated DEREs, of which the majority were Ty3 or unknown REs. On the other hand, we did not detect any significantly differentially expressed REs in the adult vs sub-adult blastemas comparison.

**Fig 4.**
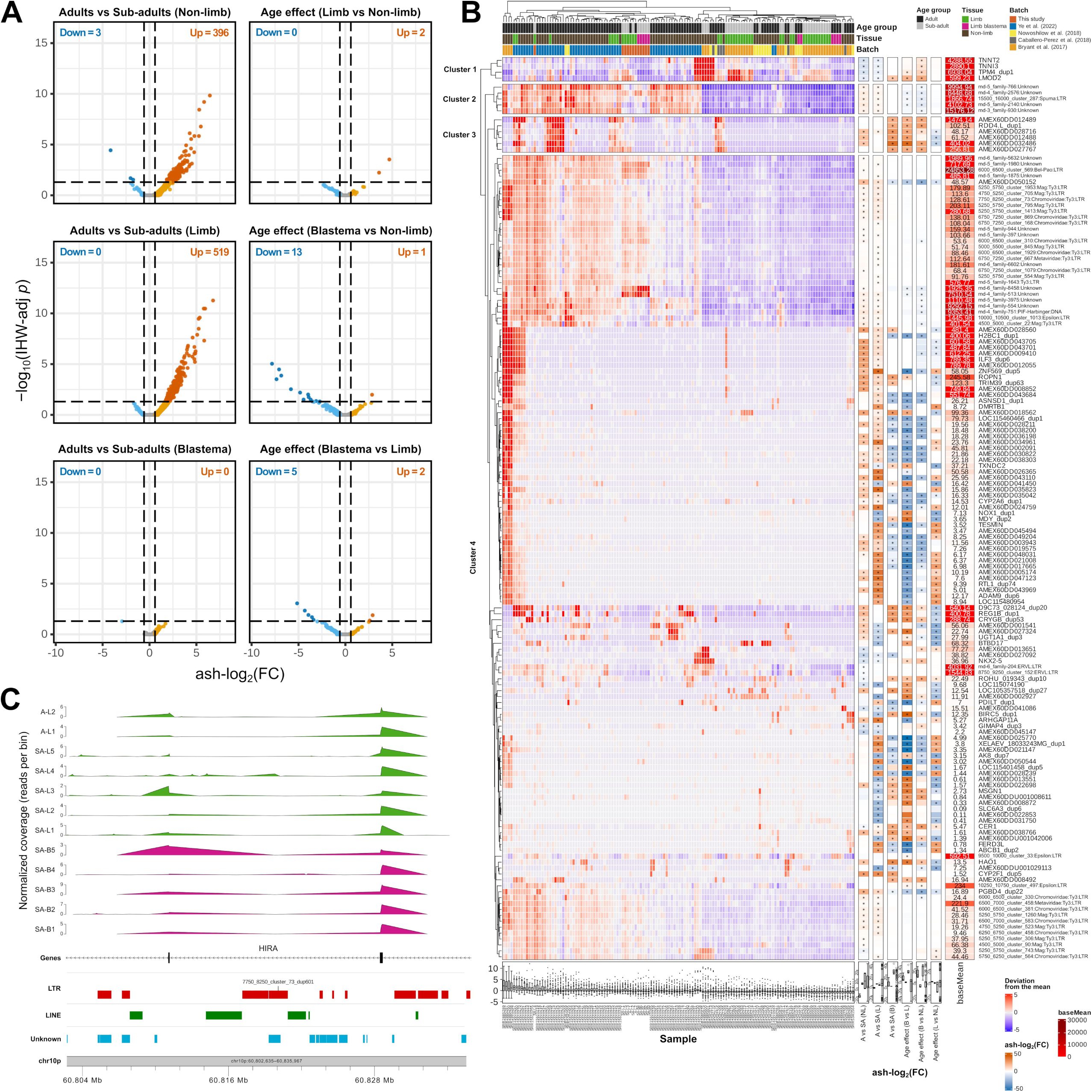
Effects of chronological aging on the repetitive element expression changes in the axolotl’s regenerating limb. (A) Volcano plots show the repetitive element (RE) expression differences between adults and sub-adults for each tissue type and the age effect differences in each pairwise comparison of tissues. Repetitive elements with significantly increased (ash-log_2_(FC) > log_2_(1.5), IHW-adj *p* < 0.05) or decreased (ash-log_2_(FC) < −log_2_(1.5), IHW-adj *p* < 0.05) expression (DEREs) are represented by dark orange and dark blue points, respectively. **(B)** Heatmap of the deviation from the mean of each VST-normalized gene/RE count, split into clusters by the global dendrogram. Columns with color-coded ash-log_2_(FC) by contrast (∗: IHW-adj *p* < 0.05) and baseMean values are shown on the right for each gene/RE. **(C)** Karyoplot of 7750_8250_cluster_73_dup601 (Chromoviridae:Ty3:LTR), one of the upregulated RE loci in the adults vs sub-adults (limb) contrast which also overlaps with an intron of *HIRA*, an upregulated gene for the same contrast. RNA-seq read coverage of each sample from this study is plotted on top. SA: sub-adult, A: adult, L: limb, B: limb blastema.

**Table 2.**
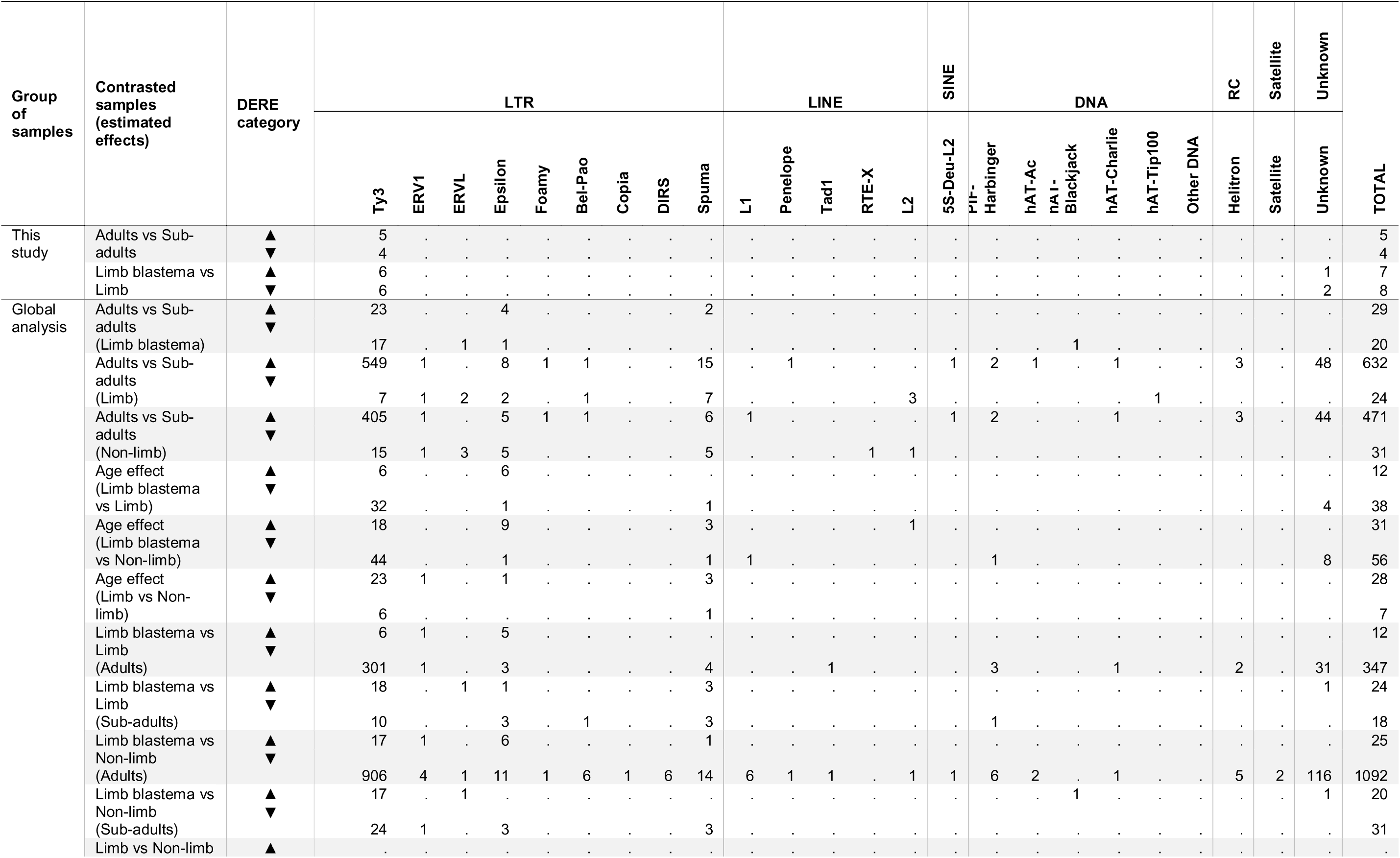
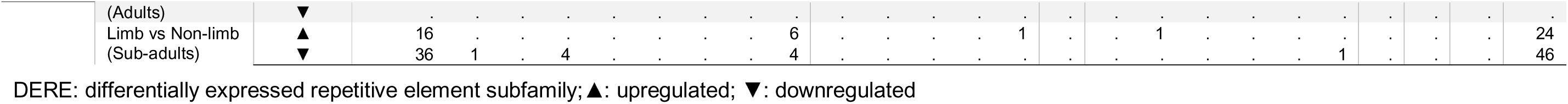
Count and annotation of significantly differentially expressed repetitive element subfamilies across contrasts of age and tissue of axolotl samples.

Consistent with what we observed in the age contrasts, the tissue comparisons revealed that during regeneration of the limb the blastema undergoes a major underexpression of mostly Ty3 retrotransposons of the Chromoviridae and Mag families, relative to samples of the limb and non-limb tissues (**S3A Fig**). This downregulation of RE expression between the blastema and the limb and non-limb tissues appears stronger in samples of adult (older) axolotls than in samples from sub-adult (younger) axolotls. Moreover, 209 (∼40 %) of the 519 repetitive elements that were upregulated in the comparison between adults and sub-adult limbs were also downregulated in the adult blastema, which represents a ∼72 % of the 304 total REs downregulated in adult blastemas when contrasted with adult limbs (**Figs 4A and S3A**). However, contrary to what we found for protein-coding genes, the difference in the effect of age on the blastema, compared to the effect it has on the limb is significant for only a few elements, including two upregulated Epsilon LTRs, one downregulated Spuma LTR, and four downregulated unknown REs (**Figs 4A**).

To analyze gene and RE expression together, we performed hierarchical clustering on the deviation from the mean of each element and then split the selected genes and REs into four clusters by the global dendrogram.

In clusters 2 and 4 of the first dendrogram (**Figs 4B**), we found a set of covarying genes/REs affected by chronological aging in the limb that makes it possible to associate several upregulated genes (e.g. *ZNF569*, *DMRTB1*, *TESMIN*, *RTL1*, *ADAM9*, *ARHGAP11A*, *PGBD4*, *H2BC1*, *FERD3L*, etc.) and LTRs (e.g. Chromoviridae, Mag, Metaviridae, Epsilon, Spuma, ERVL), with various downregulated genes (e.g. *NOX1*, *MDY*, *UGT1A1*, *BTBD17*, *BIRC5*, *GIMAP4*, *AK8*, *MSGN1*, *SLC6A3*, *TXNC2*, etc.). Additionally, there are many unannotated genes and unknown REs with a similar variation profile associated with the aged limb. In contrast, for most of the genes/REs in this set, the increase in age has a significantly different effect in the blastema. For instance, for some genes aging has the opposite effect: genes such as *PGBD4*, *FERD3L*, *TESMIN* and *H2BC1* are downregulated, while genes such as *MSGN1*, *BIRC5*, *MDY*, and *TXNC2* are upregulated. For other genes/REs, their expression is not affected by aging in the blastema: *SLC6A3*, *BTBD17*, *UGT1A1*, *ADAM9*, *RTL1*, *NOX1*, *DMRTB1*, *ZNF569*, a Spuma LTR and other unknown REs. In other cases, the effect of aging has the same direction but significantly lower (e.g., *AK8*) or higher (e.g. *CER1*) magnitude in the blastema. Finally, other genes such as *REG1B*, *CRYGB*, *ROHU_019343*, and a putative transposon DNA polymerase (*D9C73_028124*), are only affected by age in the blastema, and not in the limb.

Additionally, in the second dendrogram we identified sets of genes/REs that covary in blastemas when compared to limb tissues (**S3B Fig**). One set consists of genes with a distinct upregulation in the sub-adult blastema, including *BHLHA9*, *DDX4*, *XELAEV_18033243MG*, *PARPI_0030888*, *OVOL3*, *CCDC166*, *UCP3*, *MUC5B*, *C1ORF11*, and several unannotated genes (e.g., *AMEX60DDU001002634*, *AMEX60DD050544*, *AMEX60DD029513*). These genes showed no dysregulation in the blastema of adult axolotls. On the other hand, genes such as *CDKL3*, *RGSL2*, *OR8H3*, *ACRBP*, *CA13*, *MORC4*, *FSIP2*, *AQP8*, *CYP2A6*, and *S100PBP*, form a covarying group clearly downregulated in the regenerative tissue of sub-adult axolotls, and even more downregulated in the blastema of adults. These results suggest that the gene regulation needed during regeneration of the limb can be affected in multiple directions by an increase in age. In some cases, blastemas of aged axolotls might be unable to positively or negatively regulate the expression of regeneration-associated genes. In contrast, DEREs in these clusters exhibit a less notable dysregulation than the protein-coding genes and form a distinct set of Ty3 retrotransposons (Chromoviridae, Mag, Metaviridae) that are mostly downregulated in the blastema of adult axolotls, with only a few underexpressed in sub-adult blastemas. Among the top DEREs, only two show upregulation in blastemas: the previously identified unknown repeat (*rnd-6_family-8458*) in sub-adults, and an Epsilon LTR (*9500_10000_cluster_33*) in adults. These findings indicate that, compared to the coding genes, REs are mostly dysregulated in one direction (downregulated) even in the blastemas of aged axolotls. However, the effect of the upregulation of even a few REs might be more impactful due to their multiple genomic insertions.

Next, we annotated each locus of each DERE to its nearest genomic/genic feature to assess their vicinity to DEGs. Interestingly, the majority of non-unique DERE loci analyzed were located in intergenic regions (∼55 %) or within the introns (∼23 %) or 50 kb downstream regions (∼4.5 %) of non-DEGs (**S2 and S3 Tables**). Only a limited fraction of all DERE loci were located in the introns of downregulated (∼4.4 %) and upregulated (∼1.9 %) DEGs. For example, *7750_8250_cluster_73*, a Chromoviridae:Ty3:LTR subfamily significantly upregulated in the contrast of adult and sub-adult limbs is located within an intron of *HIRA*, a gene which is also overexpressed for the same contrast (**Fig 4C**). However, none of the DERE loci that colocalized with DEGs shared a deviation-from-the-mean cluster with a DEG. Overall, these results indicate that the potential regulation of gene expression by REs in both aging and regeneration is for the most part independent of their genomic proximity to the genes. In other words, for the DEREs that were detected outside and far from the DEGs, it is likely that their modulatory effect is not direct and requires the generation of intermediates or the interaction with other regulatory factors.

### A network of retrotransposition and muscle development genes is jointly suppressed in aging axolotls

Besides their direct impact on the expression of colocalized or neighboring genes, DEREs may also establish expression networks with other protein-coding genes, contributing to specific biological functions. To further explore the potential roles of REs in aging and in the development of the regenerative tissue of the limb, we examined the correlation between genes and REs of all samples using weighted gene co-expression network analysis (WGCNA). This analysis revealed 16 coexpression modules, with sizes ranging from 126 to 11203 genes/REs (**Figs 5A-C**). Next, to understand the physiologic significance of the modules, we correlated the 16 modules’ eigengenes (i.e., their first principal components) with the sample traits: age group (adults or sub-adults) or tissue (non-limb, limb, or blastema) (**Fig 5B**).

**Fig 5.**
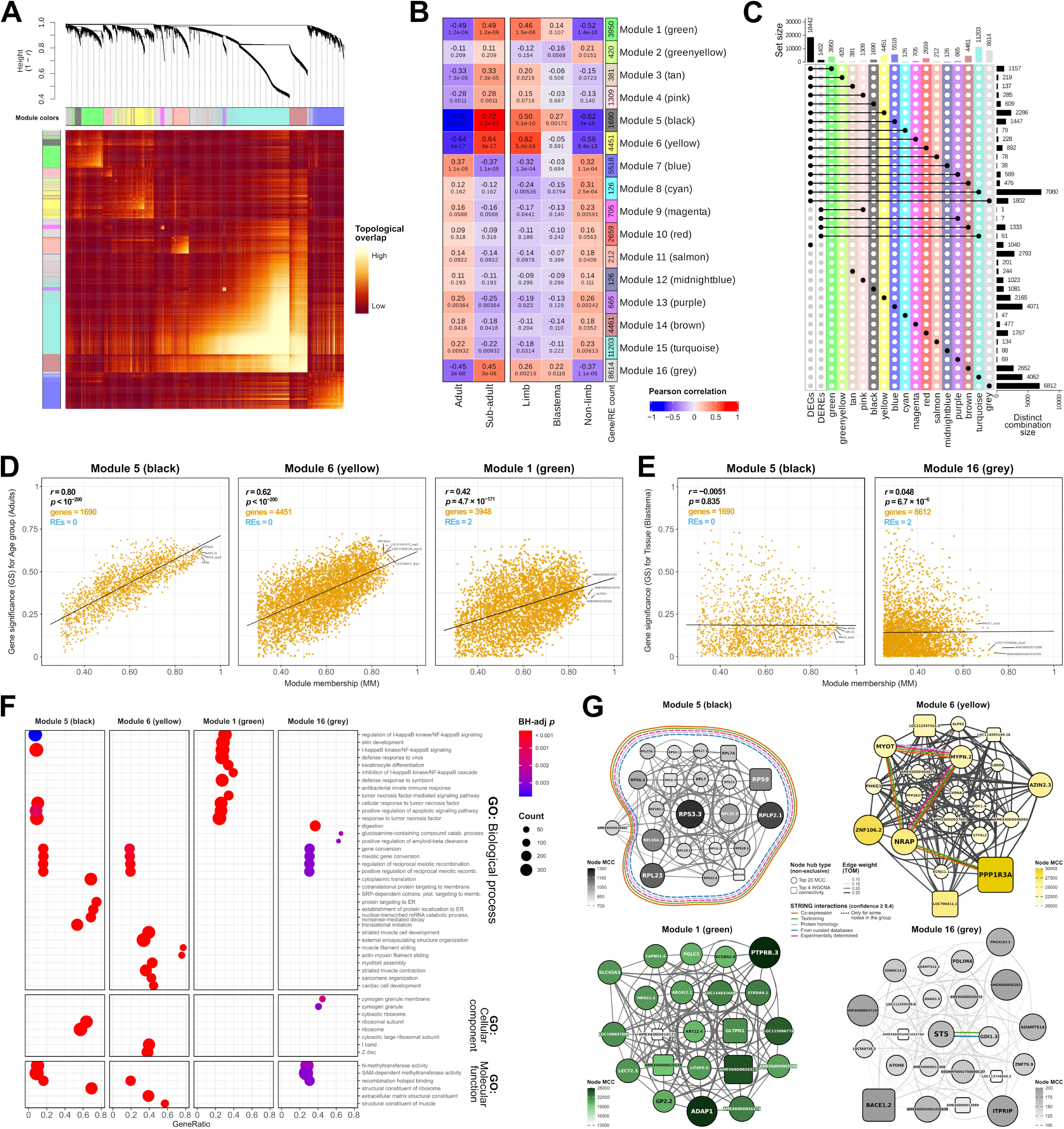
Gene and repetitive element coexpression and their association with aging and regeneration in the axolotl. (A) Heatmap plot of topological overlap in the gene and repetitive element (RE) network. Each row or column corresponds to a gene/RE, light color denotes high topological overlap, and progressively darker red denotes lower topological overlap. Lighter squares along the diagonal correspond to coexpression modules. The gene/RE dendrogram and module assignment are along the top. **(B)** Heatmap of module-trait Pearson correlation. Each cell shows the correlation value on top, and the associated BH-adj *p* below it. **(C)** UpSet plot of the intersections between module genes/REs and the complete set of differentially expressed genes and REs detected previously through differential expression analysis. **(D)** Scatterplots of gene significance (GS) for the age group trait versus module membership (MM) in the modules with highest correlation with age group. **(E)** Scatterplots of GS for the limb blastema tissue versus MM in the modules with highest correlation with this trait. **(F)** Dotplot shows the significantly enriched gene ontology (GO) terms identified for each coexpression module by overrepresentation analysis (ORA). **(G)** Coexpression subnetworks show the hub genes/REs in modules 1 (black), 6 (yellow), 1 (green) and 16 (grey) selected based on intramodular connectivity and maximal clique centrality (MCC) scores. Gene symbol numeric suffixes denote gene duplicates. Reported functional and physical protein associations are shown as color-coded edges.

The eigengenes of several modules were highly correlated with the age group trait, including module 5 (black; 1690 genes; *r* = −0.72 correlation with adult age, BH-adj *p* = 1.3 × 10^−21^), module 6 (yellow; 4451 genes; *r* = −0.64 correlation with adult age, BH-adj *p* = 7.9 × 10^−16^), module 1 (green; 3948 genes and 2 REs; *r* = −0.49 correlation with adult age, BH-adj *p* = 8.5 × 10^−9^), and module 16 (grey; 8612 genes and 2 REs; *r* = −0.45 correlation with adult age, BH-adj *p* = 1.7 × 10^−7^). We found that these modules were also highly correlated with the tissue trait, specifically the limb tissues, although only two of these modules were significantly correlated with the limb blastema trait: module 5 (*r* = 0.27, BH-adj *p* = 0.00552), and module 16 (*r* = 0.22, BH-adj *p* = 0.027). Notably, only two of these modules contain coexpressed REs: module 1 incorporates the *13750_14250_cluster_243*:Spuma:LTR and *5000_5500_cluster_187*:Mag:Ty3:LTR subfamilies, while module 16 includes the *6250_6750_cluster_443*:Chromoviridae:Ty3:LTR subfamily and an unknown RE (*rnd-3_family-511*) (**Fig 5C**). **S4 Table** presents the list of the DEGs and DEREs with the highest rate of change based on the differential expression contrasts, as well as the WGCNA module to which they belonged.

Then, to quantify the associations of individual genes/REs with our traits of interest (age group or blastema tissue), we measured their gene significance (GS), defined as the absolute value of the correlation between each gene/RE and the trait, and their module membership (MM), defined as the correlation of the module eigengene and the expression profile of the gene/RE. We found that GS and MM were highly correlated in modules 5 (black), 6 (yellow) and 1 (green), illustrating that genes/REs significantly associated with the age group of the axolotl were also the most important elements of their modules (**Fig 5D**). Although this was true for modules correlated with age group, modules associated to the blastema tissue show a minimal GS-MM correlation (**Fig 5E**), which could indicate that only a submodule relates to the blastema trait, and that this module-trait association requires further validation.

Next, we conducted overrepresentation analysis (ORA) to determine whether these modules were significantly enriched with regard to known gene ontologies (**Fig 5F**). Among the 681 significantly enriched pathways that we detected in module 5 (black), we found that the top ones were related to cytosolic ribosome components, rRNA and mRNA metabolism, translation initiation and regulation, protein targeting to membrane and ER, and liver regeneration. Module 6 (yellow) showed an overrepresentation of 1416 GO terms, mainly associated with striated muscle morphogenesis, differentiation and function (actin-myosin filament sliding, contraction regulation, etc.), as well as other development terms such as extracellular matrix structural constituents, structural molecule activity conferring elasticity, glycosaminoglycan binding, and chondrocyte differentiation. We detected a set of 1820 terms significantly enriched in module 1 (green), in which the predominant pathways were implicated in immune and inflammatory responses, cell survival and proliferation, and skin/keratinocyte development. In particular, the most significant terms in this module were involved in the negative regulation of I-kappaB kinase/NF-kappaB signaling, immune response, viral processes, and cytokine production, but also the positive regulation of the apoptotic signaling pathway. Module 16 (grey) was significantly enriched for 60 GO terms, including digestion, zymogen granule membrane, N-methyltransferase activity (mainly SAM-dependent, and H3-K4 or H3-K36 specific), meiosis-related processes (regulation, gene conversion, recombination hotspot binding, DSB repair, etc.), and positive regulation of connective tissue replacement. Subsequently, to pin down the hub elements in each module we selected the genes/REs with the highest intramodular connectivity (**Figs 5D, 5E and** 5G) and maximal clique centrality (MCC) scores (**Fig 5G**).

In agreement with the ORA results, we found that most hub genes in module 5 (black) encoded for proteins of the small and large ribosomal subunits (*RPS3*, *RPLP2*, *RPS9*, *RPL23*, etc.), as well as a eukaryotic translation elongation factor involved in the transfer of aminoacylated tRNAs to the ribosome (*EEF1B2*), and the unannotated gene *AMEX60DD030462* (**Fig 5G**). In addition, we annotated the functional and physical interactions of the hub genes based on the STRING database and observed that all proteins in the module except for the unannotated gene have reported interactions (co-expression, from text mining, curated databases, etc.).

In module 6 (yellow), the gene with the highest MCC score was *PPP1R3A*, a regulatory glycogen-targeting subunit of protein phosphatase-1 (PP1) which is essential for cell division, regulation of glycogen metabolism, muscle contractility and protein synthesis (**Fig 5G**). Other hub genes in this module were also related to muscle development and activity (*NRAP*, *ZNF106*, *MYOT*, etc.), agreeing with ORA results, and *PPP1R3A*, *MYOT*, *MYPN* and *NRAP* have previously been reported to be coexpressed. Notably, we also identified two previously reported interactors, ornithine decarboxylase 1 (*OCD1*) and its enzymatically inactive homolog antizyme inhibitor 2 (*AZIN2*) as hub genes in this module. Moreover, although we did not detect any coexpressed RE as such in module 6, we found three hub genes that encode for (predicted) protein domains characteristic of retrotransposable elements. Specifically, *LOC790411* contains LINE-1 endonuclease reverse transcriptase and transposase domains, *LOC114595149* contains an exonuclease-endonuclease-phosphatase (EEP) and a non-LTR retrotransposon/retrovirus reverse transcriptase (RT), and *LOC112547415* contains a DIRS-1 RT domain and a DIRS-1 ribonuclease HI. These findings indicate that several retrotransposons might serve as hubs and indirectly participate in the gene co-expression network of muscle development regulation and function that is suppressed by aging in the axolotl.

Based on MCC, the top hub gene in module 1 (green) was protein tyrosine phosphatase receptor type B (*PTPRB*), which plays an important role in blood vessel remodeling during embryonic development, maintenance, and angiogenesis. PTBs also regulate cell growth, differentiation, mitotic cycle, and oncogenic transformation. Interestingly, *PTPRB* is essential for the adhesive function of vascular epithelium-cadherin CDH5 in endothelial cells, and we identified another top hub gene in this module (*LOC115096778*) that encodes for a cadherin-like protein involved in cell morphogenesis, and calcium-dependent cell-cell adhesion via plasma membrane molecules. In general, most hub genes in module 1 were related to cell signaling (*ADAP1*, *ST8SIA6*), vesicle trafficking, integral composition of membranes (*PQLC3*, *LOC114632468*), cell-pathogen interaction and chemotaxis (*GP2*, *LECT2*), chondrocytes and osteoblasts (*LECT2*), among other unannotated functions (*AMEX60DD026325*) (**Fig 5G**).

Beta-secretase 1 (*BACE1*), a proteolytic processor of the amyloid precursor protein (APP), was the top hub gene in module 16 (grey), which showed a significant positive correlation with the regenerative tissue. We also found a high interaction with *PRICKLE2*, a neural development gene which has also been involved in the inhibition of WNT/PCP/JNK signaling activated by APP. Other unannotated genes also showed a high MCC score (*AMEX60DD037326*, *AMEX60DD052921*) or connectivity (*AMEX60DD013099*). Moreover, we identified other hubs involved in intracellular calcium signaling (*ITPRIP*), formation of collagen fibers (*ADAMTS14*), and regulation of MAPK1/ERK2 kinase activity (*ST5*). Interestingly, we also detected hub genes with LTR and non-LTR protein domains, and a potential regulator of their RNA pol II-specific transcription (*ZNF79*). Specifically, *LOC113746826* contains retropepsin, LTR RT, Ty3 RNase HI, IS481 transposase, and integrase domains, and *LOC568735* comprises EEP and non-LTR RT domains (**Fig 5G**). Together with our ORA results, these findings suggest that retrotransposons and transcription factors might serve as hubs and indirectly participate in the networks of connective tissue replacement, amyloid proteolysis, and N-methyltransferase pathways that are activated during regeneration in the axolotl.

## Discussion

Multiple studies have identified protein-coding genes with a role in regeneration of the limb in *Ambystoma mexicanum*. Here we have shown that there is a pattern of predominant gene downregulation between 10 dpa blastemas and limb tissues of axolotls, for the most part represented by muscle development suppression, which corroborates previous studies [24–26].

We additionally identified a significant upregulation of genes which have been previously implicated in limb regeneration in the axolotl. For instance, interleukin 11 (*IL11*) exhibited significant upregulation in the regenerative tissue, possibly related to its role in the inflammatory response [27]. It may also be involved in cellular reprogramming or act as a limiting factor for fibrotic scarring during tissue regeneration, as reported in *Danio rerio* [28]. However, our results show that *IL11* has a role at 10 dpa, meaning it is differentially expressed beyond 3-5 dpa, as described previously [26]. Another gene that we detected significantly overexpressed in the regenerative tissue, *ODAM*, has been identified as a marker in fibroblasts on trajectories towards osteoblast-like cells of the medium-bud blastema [29], and in various cancer cells [30]. In addition, we report that two understudied adapter proteins, SH2 domain-containing adapter protein D (*SHD*) and F (*SHF*) are also upregulated in the 10 dpa blastema. Based on a study implicating SHF as a regulator of apoptosis in mouse fibroblasts [31], and the reported role of apoptosis and bone remodeling processes during regeneration [32], these adapter proteins serve as a promising focal point for further research in the axolotl. Interestingly, we found that basic helix-loop-helix family member A9 (*BHLHA9*), which encodes a transcription factor involved in the regulation of apoptosis during autopod development [33] is also significantly upregulated in the limb blastema. Briefly, BHLHA9 has been reported to regulate apical ectodermal ridge (AER) formation during mouse limb/finger development by regulating the expression of genes such as *TP63* and *FGF8*, which serve as key molecular modulators of apoptosis and limb formation [34]. To our knowledge, BHLHA9 does not have a reported role in the axolotl blastema and is therefore a promising candidate for future investigations into the apoptotic process during limb regeneration in *A. mexicanum*.

Even though there is a growing recognition of the role of TEs in development, their contribution to animal regeneration has just recently started to receive attention [3]. Repetitive element transcriptional activity and differential expression during regeneration has only been studied in a few animal models, including *Drosophila* [10], the brown rock sea cucumber *Holothuria glaberrima* [8] and *A. mexicanum* [9]. In the case of the axolotl, Zhu *et al.* [9] described both overexpression and higher retrotransposition of LINE-1 in dedifferentiating tissues of the blastema. While they suggest that the reactivation of the LINE-1 retrotransposon may serve as a marker for cellular dedifferentiation in the early-stage of limb regeneration, they do not suggest a precise function for it, nor do they describe the expression of other elements in regeneration. Here, we characterized the expression profiles of all currently modeled REs in the regenerating limb of *A. mexicanum*. We showed that in the blastemas (10 dpa) of native (Xochimilco) axolotls, only 3 Mag and 3 Chromoviridae retrotransposon (Ty3 superfamily), and 1 unannotated repeat exhibit signs of significant upregulation. Additionally, we report that only 3 Mag and 3 Chromoviridae retrotransposons, and 2 unannotated repeats are downregulated in the native blastema. Interestingly, in contrast with the results of Zhu *et al.* [9], we did not detect significant differential expression of any subfamily of LINE-1 in the blastema of native axolotls, but we did in the context of aging.

By incorporating axolotl tissue samples from multiple studies, we also showed that during regeneration the blastema undergoes a major underexpression of mostly Ty3 retrotransposons of the Chromoviridae and Mag families relative to the limb samples. In fact, we detected only 2 upregulated REs, an Epsilon LTR in adults (*9500_10000_cluster_33*), and an unknown element in sub-adults (*rnd-6_family-8458*). These findings conform with the idea that differential expression of retroelements in regeneration is a large-scale non-specific phenomenon [3], in other words, that REs are regulated *en masse* after an injury and upon the development of the regenerate.

Nonetheless, our observation of a much more predominant downregulation of REs is not concordant with the traditional view of RE dysregulation during regeneration, which poses that transposons only make use of the global injury-induced chromatin activation and transcriptional de-repression (i.e., an overall reduction of epigenetic silencing that leads to a germline-like state) to amplify themselves and thereby increase their odds of surviving the lysis of their host cell [3,9]. On the contrary, the obvious suppression of Ty3 transposons we describe here suggests a much more finely tuned epigenetic regulation of transposon expression during limb regeneration. Conceivably, TE silencing in the blastema might be achieved through DNA methylation [35] or Polycomb-associated histone modification redistribution [36]. These mechanisms could also exert its repressive effects on TE transcription by sterically interfering with activator transcription factors (TF) binding sites in TE promoter proximal regulatory regions [35], or by increasing preferential binding of TE-silencing TFs such as C2H2-ZNFs [37] to methylated CpGs within TE promoters [38]. However, further studies are still needed to pinpoint the specific function of each subfamily of repetitive elements and the epigenetic mechanisms responsible for their silencing during limb regeneration in the axolotl.

Several studies in multiple models have reported that repetitive elements accumulate progressively with age. For instance, studies in mice, flies, and other organisms have shown that active or transposable RE can contribute to the aging process [13,14], and RE transcript accumulation is involved in age-related neurodegenerative diseases [15]. Also, increases in RE transcript expression have been used to accurately predict biological age in human fibroblasts and *C. elegans* samples [19]. Until now, there has been no research into how chronological aging affects RE expression in *Ambystoma mexicanum*. The analyses we present here demonstrate a pattern comparable to that reported for other species, in which the limbs and other tissues of the axolotl undergo a significant age-related RE upregulation, mostly dominated by LTR retrotransposons of the Ty3 superfamily.

A profile of Ty3 upregulation such as the one we report here for aging axolotls can reflect a higher transcription by the host’s RNA polymerase II (an overexpression of TE mRNAs) or an increase in the number of genomic insertions of the TE cDNA, which would also then increase the rate of TE transcription events in the genome. Studies in *Drosophila* have revealed that endogenous overexpression of TEs during aging does not necessarily lead to new genomic insertions [14]. Moreover, reducing the expression of two different Ty3 retrotransposons (*412* and *Roo*) extends lifespan without increasing TE mobilization, suggesting that the presence of TE mRNAs or even DNA intermediates, and not insertion, is the detrimental factor to cell and animal function [14]. This could also be the case for the age-related TE upregulation observed in the axolotl, however, these two scenarios are indistinguishable by means of the RNA analyses we present in this study.

Retrotransposon activity can regulate gene expression and introduce both beneficial and catastrophic events to the host genome. The mechanisms by which TEs are able to affect gene-regulatory networks include the addition of TF binding sites, promoters, enhancers, and splicing sites, among others, but TE-derived sequences can also undergo co-option/exaptation/domestication as effector proteins [39]. These processes can lead to the gene expression changes that underlie some diseases [5]. Our comparison of native adult axolotls against their younger counterparts revealed a potential aging-related gene/TE upregulated cluster involving a LINE-1 transcript (*ORF2*), a non-LTR retrotransposon (*LOC115370421*), a tumor suppressor and immune detector of viral repeats (*DMBT1*), a suppressor of DNA and histone methylation overexpressed in human cancers (*NNMT*), and two DNA-binding transcription factors (*ZNF665* and *ZNF501*). Among these, ORF2 is particularly interesting because it provides the reverse transcriptase activity required for target-primed reverse transcription of the LINE-1 mRNA, a crucial step in its retrotransposition [40]. Additionally, the KRAB-zinc finger protein ZNF665 is also noteworthy, since previous work has revealed that many Krüppel-associated box domain (KRAB) zinc finger proteins (KZFPs) recognize TE-embedded sequences as genomic targets, and that KZFPs suppress TEs or facilitate the co-option of their regulatory potential for the benefit of the host [37]. In fact, in humans, ZNF665 has targets of the LINE-1/L1 family, specifically, L1M3f and L1MA4 elements [37]. Overall, the joint upregulation of these genes could mean that an age-induced overexpression of *NNMT* might cause a reduction of the methylation levels in KZFPs binding sites within LINE-1/TE promoters, thus reducing the affinity for KZFPs and their repressive potential over TEs. This would potentially generate an upregulation of KZFPs to counteract TE overexpression. Furthermore, NNMT could also affect the activity of the DMBT1 protein through hypomethylation, as reported for other tumor suppressors such as PP2A/PPP2CA [41].

Besides the Ty3 LTR predominant upregulation caused or affected by age, we have also shown that a number of Ty3 and other non-LTR retrotransposons are downregulated by aging. For instance, some loci with predicted non-LTR retrotransposon domains (*LOC790411*: LINE-1 RTs and transposases, and *LOC112547415*: DIRS-1 RTs and ribonucleases) act as secondary hubs of a muscle development gene expression network that is jointly downregulated by aging in the axolotl. The main interactor in this network is PPP1R3A, the regulatory subunit of a glycogen-associated form of protein phosphatase-1 (PP1) that mediates muscle contraction and synthesis. Interestingly, we found also several downregulated non-KRAB ZNFs (e.g., *ZNF106*) and KZFPs (e.g., *ZNF133*, *ZNF160*, *ZNF182*) in the network, suggesting an integrated regulation of non-LTR retrotransposons and PP1-related functions by TFs. Evolutionarily recent KZFPs almost universally target TEs, often have paralogs, and display a protein interactome that represses transcription by forming a heterochromatin-inducing complex based on TRIM28 [37], another PP1 interactor involved in muscle function. The analysis of this expression network, hints at an interaction between three PP1-related proteins (PPP1R3A, PPP1R27, TRIM28/PPP1R157), ZNF transcription factors (ZNF106, ZNF133, etc.) and non-LTR retrotransposon expression.

With these data in mind, we suggest that a major factor in the suppression of skeletal muscle development and function caused by aging in the axolotl is the downregulation of *PPP1R3A* expression and, with it, the limitation of its glycogen binding activity to PP1. Considering that in humans 95 % of the promoter sequence of *PPP1R3A* is derived from LTR-TE sequences [42], and since KZFPs are able to recognize TE-embedded sequences and facilitate their regulatory potential [37], it could be hypothesized that the underexpression of several KZFP transcription factors has a direct detrimental effect on the expression and activity of PPP1R3A. Importantly, ZNF106 is a C2H2-ZNF but does not contain a KRAB domain, which is usually required for TRIM28-mediated transcription regulation [43]. However, other suppressed KZFPs in the same network (e.g., *ZNF133*, *ZNF160*, *ZNF182*, etc.) could have similar affinity for TE-derived promoters of other muscle development genes, or for the other hub LINE-1 and DIRS-1 retrotransposons that we found downregulated in the module. Still, further studies in the axolotl are needed to establish the origin and RE composition of the *PPP1R3A* promoter sequence and its potential as a TF binding site for C2H2-ZNFs. Furthermore, it must be determined if and which KZFPs function as activators or repressors of TEs or TE-derived cis-regulatory sequences during age-related muscle suppression, and whether TRIM28, PPP1R3A, PPP1R27 or other PP1 regulatory subunits direct this protein interactome.

In general, in both chronological aging and regeneration, we found a significant dysregulation of numerous C2H2-ZNFs and KZFPs coupled with differentially expressed LTR and non-LTR retrotransposons, also forming coexpression networks. Furthermore, we also identified special cases in which single genes, highly dysregulated by regeneration or aging, contained both LTR transposition enzymes and C2H2-ZNF domains (e.g., *LOC105357518*). Although in most cases the changes in TF and TE expression occurred in the same direction, there were instances with opposing regulation, suggesting intricate mechanisms or external factors influencing TFs and TEs independently. In brief, we propose that KZFPs and other C2H2-ZNFs may be important regulators of the retrotransposon expression during aging and regeneration in the axolotl. Of note, the potential of TFs, specifically KRAB-ZFPs, to play a major role in TE activity regulation during tissue regeneration has recently been hypothesized by Angileri *et al.* [44]. Even though most KZFPs have been reported to be repressive of TEs in other organisms, the direction of their regulatory activity in this context might be dependent on the specific KZFP and TE. Moreover, DNA methylation could be one of the main epigenetic mechanisms that fine-tune the transcription of TEs or genes with TE-derived promoters/enhancers during aging and also during blastema development.

We previously hypothesized that physiological or pathological processes such as aging may alter the expression of repetitive elements and consequently modify the basal state from which the axolotl responds to tissue damage and regulates regeneration. In favor of this premise, we have shown that more retrotransposons and protein-coding genes are downregulated in the blastema of aged axolotls than in the blastema of younger axolotls. Moreover, ∼40 % of the retrotransposons and ∼54 % of the protein-coding genes whose expression was upregulated by aging in the limb, were then downregulated in the blastema. This means that, while there is a significant counterresponse during the development of the regenerate to previous age-related gene and RE dysregulation, there are still several genes and REs that remain affected and may impact future regenerative processes. Retrotransposons might shift the immune response or metabolism, that potentially lead to the age-related decrease in regenerative capacity in the axolotl not only by direct gene regulation (e.g., through ncRNAs or genomic insertions), but also by producing retrotransposition intermediates. In fact, because of their similarity to retroviruses, cytosolic DNA intermediates produced by reverse transcriptases or even the enzymes themselves could trigger the activity of the immune system [14], induce the nuclear factor kappa B (NFκB) pathway or the interferon beta (IFNβ) inflammatory response [13,45], and lead to chronic inflammation and cellular senescence [45], affecting the ability of *A. mexicanum* to regenerate a limb.

In summary, in this study we provide a collection of genes and repetitive elements in the axolotl for which we have shown significant dysregulation as an effect of chronological aging or throughout the development of the blastema of the limb. Moreover, we reveal that repetitive elements, mostly of the Ty3 superfamily, are predominantly upregulated as an effect of aging in the limb of *A. mexicanum*. In contrast, during regeneration of the limb, Ty3 retrotransposon expression is largely downregulated. Although most differentially expressed retrotransposons did not have a genomic overlap with differentially expressed genes, we identified several coexpression modules with both protein-coding genes and LTRs/non-LTRs, suggesting an indirect TE-mediated regulatory network directing processes such as a general muscle development suppression. We highlighted that although the blastema is able to readjust most of the transposon dysregulation caused by aging, there are still several transposons that remain affected and may impact the immune response or metabolism in the regenerative process. We identified numerous C2H2-ZFPs coexpressed with hub retrotransposons and emphasized their potential as regulators of transposable element expression during aging and regeneration in the axolotl. Overall, we argue that RE expression changes during reparative regeneration in the axolotl do not represent a simple exploitation of the transcriptional machinery of the host, but rather a finely tuned response to the physiological state before and after an injury, specifically controlled at the tissue and repetitive element levels.

### Limitations of the study

Even though we have shown that repetitive elements exhibit a clear pattern of predominant upregulation as an effect of chronological aging in the axolotl, further transcriptomic studies with specimens of more precise age groups (e.g., one group per year of age) could provide the means to define if RE transcript levels are a good marker of biological age in *A. mexicanum* [19], as well as to reveal the intermediate dynamics and physiological effects of RE expression through aging. Also, in the context of regeneration, analysis of RE expression in blastemas of diverse organs at more hourly or daily intervals after injury may facilitate the determination of the mechanisms by which retrotransposons regulate gene expression, metabolism, or immune response at the different stages of tissue formation.

Furthermore, in the present study we integrated RNA-seq samples from native (Xochimilco) axolotls and specimens of various laboratory strains. This allows for generalization at the species level but overlooks the importance of the genotype of the axolotl on its physiology or regenerative process. For instance, there has been phenotypic evidence indicating that neither aging nor repetitive amputations have an important effect on the regenerative potential of urodeles, while other studies argue they can be detrimental [reviewed in 46]. We have documented here and in a previous study [47] that two 8-year-old Xochimilco axolotls did not regenerate the hindlimb even after their first amputation. This opposing evidence may be due to the strain under study: impairment of regeneration after repeated amputations has been reported for American strains, and not for European strains [46]. Moreover, the hybridization and prolonged breeding history of the main laboratory strain used today may have selected against developmental or metabolic processes involved in regeneration, as reported for spontaneous metamorphosis [48,49]. Overall, these reports highlight the importance of taking the strain and genotype into account for aging and regeneration analyses in the axolotl and stress the need for advancing genomic and transcriptomic studies in native Xochimilco axolotls.

## Materials and Methods

### Sample collection

Sample extraction and RNA sequencing were conducted as described previously [47]. All biological samples were obtained from a captive population of Xochimilco Mexican axolotls (*Ambystoma mexicanum*) residing at the Unidad de Manejo Ambiental (UMA), Centro de Investigaciones Biológicas y Acuícolas de Cuemanco (CIBAC), Universidad Autónoma Metropolitana - Unidad Xochimilco (UAM-X), Mexico City, Mexico. This captive population of *A. mexicanum* at the UMA, CIBAC was established in 2007 from 31 founder organisms of wild origin (from their habitat in Xochimilco) and careful reproduction has been conducted to prevent endogamy or inbreeding.

Sedation of the axolotls was accomplished through a 20-minute immersion in a tank containing benzocaine at a concentration of 50 mg/L prior to the amputation process. Biological samples were meticulously collected, using a stereoscopic microscope, from the metatarsi of the (posterior) limbs of five male wild-type sub-adult axolotls (aged 8 months), as well as from two male wild-type adult axolotls (aged 8 years). Ten days after the amputation, blastema tissues were collected only from the five sub-adult axolotls, since none of the adult axolotls displayed development of blastema tissue even after 6 months of monitoring. In total, 12 tissue samples were collected. No animals were sacrificed for the purpose of this study, and all the axolotls were safely reintroduced to their habitat at the UMA, CIBAC following the final sample collection.

### RNA extraction and sequencing

The collected tissues were preserved in RNAlater (Invitrogen), at 4 °C for a maximum of 24 hours prior to processing. The RNA was extracted according to the protocol established by Peña-Llopis and Brugarolas [50]. RNA sample quality was assessed by High Sensitivity RNA Tapestation (Agilent Technologies Inc., California, USA) and quantified by Qubit 2.0 RNA HS assay (ThermoFisher, Massachusetts, USA). Ribosomal RNA depletion was performed using the RiboZero Gold kit (Illumina, California, USA). The rRNA-depleted RNA was purified by 2x RNAClean XP beads (Beckman Coulter) and eluted in nuclease-free water. Library construction was performed based on the manufacturer’s recommendation for SmarterStranded V2 kit (Takara Bio, California, USA). The final library quantity was measured by KAPA SYBR FAST qPCR and library quality was evaluated by TapeStation D1000 ScreenTape (Agilent Technologies, California, USA). Illumina 8-nt dual-indices were used. Equimolar pooling of libraries was performed based on QC values. Paired-end (PE) sequencing was carried out on an Illumina HiSeq sequencer (Illumina, California, USA) with a read length configuration of 150 bp and 20 million reads per sample (10 million reads in each direction).

### External RNA-seq datasets

In addition to the 12 RNA-seq datasets from native axolotl samples collected in this study, 124 datasets from previously published studies were also analyzed [4,21–23]. Each dataset was assigned a tissue group (non-limb, limb, or limb blastema) and an age group (adult or sub-adult) based on its available metadata. For samples for which there was no described age group between these two, the age group was assigned based on the age of sexual maturation reported for the axolotl [2]. Male axolotls 9 months old or younger and females 12 months old or younger were assigned as sub-adults; older axolotls were determined to be adults. The metadata of these datasets are presented in **S1 Table**.

### Genomic element annotation

The axolotl reference genome (AmexG_v6.0-DD), transcriptome (AmexT_v47-AmexG_v6.0-DD), proteome (AmexT_v47_protein) and their annotations were downloaded from https://www.axolotl-omics.org/assemblies [4].

Protein-coding sequences from the proteome file were functionally annotated using eggNOG-mapper 2.1.12 based on eggNOG 5.0 orthology data. Sequence searches were performed using DIAMOND 2.1.8 in ultra-sensitive mode. Queries were realigned with HMMER 3.3.2 to the PFAM domains found on the orthologous groups and a list of their positions on the queries was reported.

Transcripts in the transcriptome annotation (GTF) file were functionally re-annotated for GO terms, KEGG Pathways, and NCBI/Entrez Gene IDs by querying their *transcript_name(s)* against the genome-wide annotations of human, mouse, and *Xenopus*. Those *transcript_name(s)* with the [hs] (*Homo sapiens*) identifier were queried against org.Hs.eg.db, and those with [nr] (non-redundant) or no identifier were queried against org.Hs.eg.db, org.Mm.eg.db, and org.Xl.eg.db. All queries for each transcript were merged, duplicates were removed, and each transcript was assigned a comprehensive set of GO IDs, KEGG PATHWAY IDs and NCBI Gene IDs. Transcript annotation was grouped by *gene_id*, duplicates were removed, and each gene was assigned a set of GO IDs and KEGG PATHWAY IDs. Each gene was assigned a *gene_name* (symbol) by selecting among the list of *transcript_name(s)* for the gene, with the following priority: 1) first [hs] *transcript_name*, 2) first *transcript_name* annotated by eggNOG-mapper, 3) first [nr] *transcript_name*, 4) first [no identifier] *transcript_name*. In cases where no *transcript_name* could be annotated for a gene (the field was either empty, “UNKNOWN”, “N/A”, or not available), the *gene_id* was copied to the *gene_name*.

The reference RE library (AmexG_v6.0-DD_Repeats) was retrieved from the UCSC Genome Browser at https://genome.axolotl-omics.org using the UCSC Table Browser tool to query the Annotated Repeats track data hub: https://www.axolotl-omics.org/trackhubs/AmexG_v6.0-DD/Repeats/hub.description [4].

### RNA-seq data analysis

The quality of the RNA-seq datasets was assessed with FastQC 0.12.1 and summarized with MultiQC 1.19.

Gene and RE expression were quantified following two independent approaches. In the first approach, all the RNA-seq datasets were aligned to the axolotl reference genome using STAR 2.7.11a with parameters *--outFilterMultimapNmax 200 --winAnchorMultimapNmax 400 -- outFilterMismatchNoverLmax 0.04 --outFilterType BySJout*. Gene and RE family expression was then quantified with TEcount under TEtranscripts 2.2.3 with parameters *--stranded no --mode multi* and the reference gene (AmexT_v47-AmexG_v6.0-DD.gtf) and RE (AmexG_v6.0-DD_Repeats.gtf) annotations. Quantification results for all samples were joined and imported as a count matrix. In the second approach, quantification was performed on the RNA-seq datasets separately using Salmon 1.10.2 for transcript expression and ExplorATE v0.1b for RE expression. Using tximport 1.26.0, transcript and RE locus abundances were imported and summarized to the gene and RE subfamily levels, respectively.

#### Gene and repetitive element differential expression analysis

The imported quantification data was analyzed for differential expression using DESeq2 1.38.0 with two different designs. In the first design, only the 12 native axolotl samples were analyzed under an additive model *∼ Age_group + Tissue_group*. In the second design, both native and previously published samples were analyzed under an interaction model considering batch effect *∼ Batch + Age_group + Tissue_group + Age_group:Tissue_group*. Two-tailed threshold-based Wald tests of significance were performed using *altHypothesis = “greaterAbs”, lfcThreshold = log2(1.5)*. Independent filtering and *p-value* adjustment were performed using independent hypothesis weighting (IHW 1.26.0) with a significance cutoff *alpha = 0.05*. Logarithmic fold changes (LFCs) were shrunk using adaptive shrinkage (ash) with a *lfcThreshold = log2(1.5)*. Differentially expressed genes (DEGs) or REs (DEREs) of interest for a specific contrast were defined as those presenting *|ash-log_2_(FC)| > log_2_(1.5)* and *IHW-adj p < 0.05*. Volcano plots were generated with EnhancedVolcano 1.16.0 and personalized with ggplot2 3.4.4. Heatmaps and Upset plots were generated with ComplexHeatmap 2.14.0. Since two quantification approaches were used for both gene and RE expression, the findings regarding DEA in the Results section are presented as ranges (e. g. “we identified 2–6 downregulated genes…”).

#### Repetitive element genomic context annotation

All DERE loci coordinates were extracted from the repeat annotation file (AmexG_v6.0-DD_Repeats) and annotated to their nearest genomic/genic features as recorded in the transcriptome (AmexT_v47-AmexG_v6.0-DD) using UROPA 4.0.3 with the following priority: 1) inferred TSS-promoter (-1000 bp, +100 bp from gene’s 5’); 2) inferred TTS (-100 bp, +1000 bp from gene’s 3’); 3) coding sequence (CDS); 4) 5’-UTR; 5) 3’-UTR; 6) other exonic; 7) intronic; 8) other genic; 9) 5 kb upstream of gene; 10) 5 kb downstream of gene; 11) 10 kb upstream of gene; 12) 10 kb downstream of gene; 13) 50 kb upstream of gene; 14) 50 kb downstream of gene; 15) intergenic region.

#### Annotation of the protein domains in differentially expressed repetitive elements

The DNA sequences of all DEREs were extracted from the axolotl repeat library (AmexG_v6.0-DD_Repeats) and all loci of each subfamily were aligned with MAFFT v7.520. Columns in the multiple sequence alignments (MSAs) with 80 % or more gaps were removed with T-COFFEE 13.46.0.919e8c6b. The consensus sequences were generated with the *cons-plurality 1* function of the EMBOSS 6.6.0 package and queried against RepeatMasker’s *RepeatPeps.lib* with BLASTX 2.15.0, and against InterPro 98.0 with InterProScan 5.66. Additionally, the consensuses were translated with both *getorf-minsize 200* and *transeq* from EMBOSS, and then queried against PFAM 36.0 with *pfam_scan.pl* 1.6.

#### Gene set enrichment analysis based on differential expression

The complete gene ash-log_2_(FC) lists that resulted from each contrast of the DEA based on TEcount counts were subjected to fast gene set enrichment analysis (FGSEA) under the GSEA function of clusterProfiler 4.6.0, using parameters *exponent = 1, minGSSize = 10, maxGSSize = 500, eps = 0, pvalueCutoff = 0.05, pAdjustMethod = “BH”, nPermSimple = 1000000*.

#### Gene and repetitive element coexpression analysis

Gene and RE counts from the TEcount quantification of all samples were subjected to batch effect adjustment with ComBat-seq of sva 3.46.0, specifying age group and tissue as biological covariates to preserve their signal in the adjusted data. The adjusted counts were subjected to variance-stabilizing transformation (VST) and used as input for coexpression analysis, which was performed using the weighted correlation network analysis (WGCNA) R package v1.71. To calculate the adjacency matrix *a_ij_* of the signed network (*a_ij_* = |(1 + cor(*x_i_*,*x_j_*))/2|^β^), the co-expression similarity was raised to a soft-thresholding power of β = 8, which was chosen based on the criterion of approximate scale-free topology implemented in the *pickSoftThreshold* WGCNA function, i.e. this was the lowest power for which the scale free topology fitting index *R^2^* exceeded 0.85. Then, the *blockwiseModules* function of WGCNA was used to calculate the topological overlap matrix (TOM), construct the network, and group the genes and repetitive elements into co-expression modules in one step. The parameters entered were *power = 8, networkType = “signed”, deepSplit = 2, pamRespectsDendro = F, minModuleSize = 30, reassignThreshold = 0, mergeCutHeight = 0.25, maxBlockSize = 52000*.

The module eigengenes were then correlated with the phenotypes (age groups or tissues) using the *cor* (Pearson) function of WGCNA. The vectors of *p* values were generated using the *corPvalueStudent* function of the same package, and then fitted with the Benjamini-Hochberg method. Gene significance (GS) and module membership (MM) scores and correlations were defined and calculated as per the WGCNA paper.

#### Overrepresentation enrichment analysis for coexpression modules

The enrichment of biological pathways in each coexpression module was examined with overrepresentation analysis based on the hypergeometric test, as implemented in the *fgseaORA/fora* function of FGSEA 1.24.0. The GO annotation generated previously was used to produce a list of GO IDs and their corresponding axolotl genes to employ as reference. Each module’s enrichment was analyzed independently: the set of query genes was defined as all the genes detected by WGCNA for the module, and the gene universe/background was defined as the set of genes for which at least 3 samples of any of the traits significantly correlated to the module (BH-adj *p* < 0.05) had a raw TEcount count of at least 10. The minimal and maximal sizes of a gene set to test were set to 10 and 500 genes, respectively.

#### Hub gene and repetitive element detection

The hub genes/REs in the coexpression modules were identified using two ranking methods. Firstly, a modified version of the *chooseTopHubInEachModule* function of the WGCNA package was used to pinpoint the four genes with the highest connectivity in each module, looking at all genes in the expression file. Secondly, the topological overlap matrix of each coexpression module was entered into the *exportNetworkToCytoscape* function of the WGCNA package to export each weighted network to edge and node files. An adjacency threshold of 0.02 was used to filter the edges to output. The nodes and edges were imported to Cytoscape 3.10.1 and the top 20 hub genes/REs within each module were identified based on their maximal clique centrality (MCC) score as calculated with the cytoHubba 0.1 plug-in. The union set of both ranking methods was used to generate a new network on which the following algorithms were then applied for visualization: edge-weighted spring embedded layout (based on weight/TOM), yFiles (v1.1.3) Remove Overlaps, and yFiles (v1.1.3) Organic Edge Router. Next, the Cytoscape StringApp 2.0.2 was used to retrieve the functional and physical protein associations (with a confidence score ≥ 0.4) between the identified hub genes from the STRING database, which then were annotated on the networks manually.

## Ethics statement

Samples of axolotl tissue were extracted from the *A. mexicanum* colony at the Unidad de Manejo Ambiental (UMA), Centro de Investigaciones Biológicas y Acuícolas de Cuemanco (CIBAC), Universidad Autónoma Metropolitana - Unidad Xochimilco (UAM-X), Mexico City, Mexico. The UMA and colony were founded with the approval of the Secretaría de Medio Ambiente y Recursos Naturales (SEMARNAT), Mexico, under record DGVS-CR-IN-0952-D.F./07. The sample extraction protocol followed the ethical guidelines and approval (reference number CEI.2023.007) of the Comité de Ética en Investigación (CEI), División de Ciencias Biológicas y de la Salud (DCBS), UAM-X. All the experimental procedures were supervised by veterinarians at the UMA, CIBAC, according to the norms and regulations set by the SEMARNAT (NOM-087-SEMARNAT-SSA1-2002, NOM-059-SEMARNAT-2010, NOM-033-SAG/ZOO-2014). The project was registered with the Comisión Nacional de Bioética (CONBIOÉTICA), Secretaría de Salud, Mexico, under reference number CONBIOÉTICA-09-CEI-017-20180924.

## Availability of data and materials

The raw RNA-seq data from the native axolotl samples was deposited in NCBI’s Gene Expression Omnibus (GEO) under accession ID <u>GSE237864</u>. The external RNA-seq datasets are available under accessions <u>GSE92429</u>, <u>GSE182746</u>, <u>PRJNA354434</u> and <u>PRJNA378982</u>. All the underlying code for the bioinformatic analyses of native and external RNA-seq datasets is detailed in the following GitHub repository: https://github.com/samuelruizperez/axoloTE. The datasets resulting from these analyses are available in the following Zenodo repository: https://doi.org/10.5281/zenodo.10728952.

## Author contributions

**Conceptualization:** Samuel Ruiz-Pérez, Nicolas Alcaraz, Rodrigo González-Barrios. **Data curation:** Samuel Ruiz-Pérez. **Formal analysis:** Samuel Ruiz-Pérez. **Funding acquisition:** Rodrigo González-Barrios, Ernesto Soto-Reyes. **Investigation:** Samuel Ruiz-Pérez, Karla Torres-Arciga. **Methodology:** Samuel Ruiz-Pérez. **Project administration:** Rodrigo González-Barrios, Ernesto Soto-Reyes, Cynthia Gabriela Sámano-Salazar. **Resources:** Rodrigo González-Barrios, Clementina Castro-Hernández, Ernesto Soto-Reyes, José Antonio Ocampo-Cervantes, Alejandra Cervera. **Software**: Samuel Ruiz-Pérez. **Supervision**: Nicolas Alcaraz, Rodrigo González-Barrios, Ernesto Soto-Reyes. **Visualization:** Samuel Ruiz-Pérez. **Writing – Original Draft Preparation:** Samuel Ruiz-Pérez, Nicolas Alcaraz, Rodrigo González-Barrios. **Writing – Review & Editing:** Samuel Ruiz-Pérez, Karla Torres-Arciga, Nicolas Alcaraz, Alejandra Cervera, Ernesto Soto-Reyes, Rodrigo González-Barrios

## Acknowledgements

We thank Aylin del Moral-Morales for her support with access to additional high-performance computing at the *esrs-epigenetics* server of the Universidad Autónoma Metropolitana, Unidad Cuajimalpa.

