## Supplementary figures and images for "Retrotransposon expression is upregulated by aging and suppressed during regeneration of the limb in the axolotl (*Ambystoma mexicanum*)"

### S1 Fig

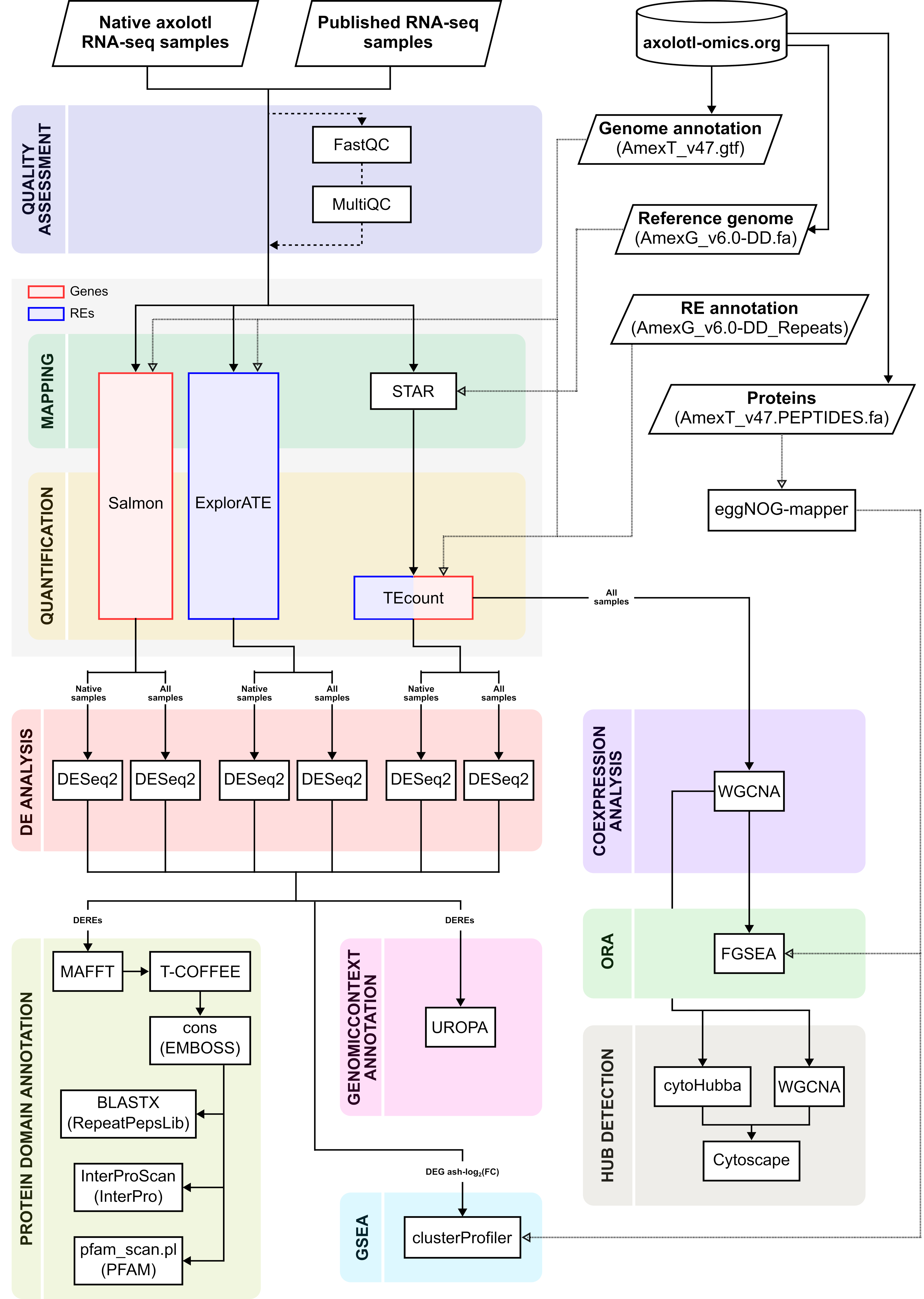

### S2 Fig

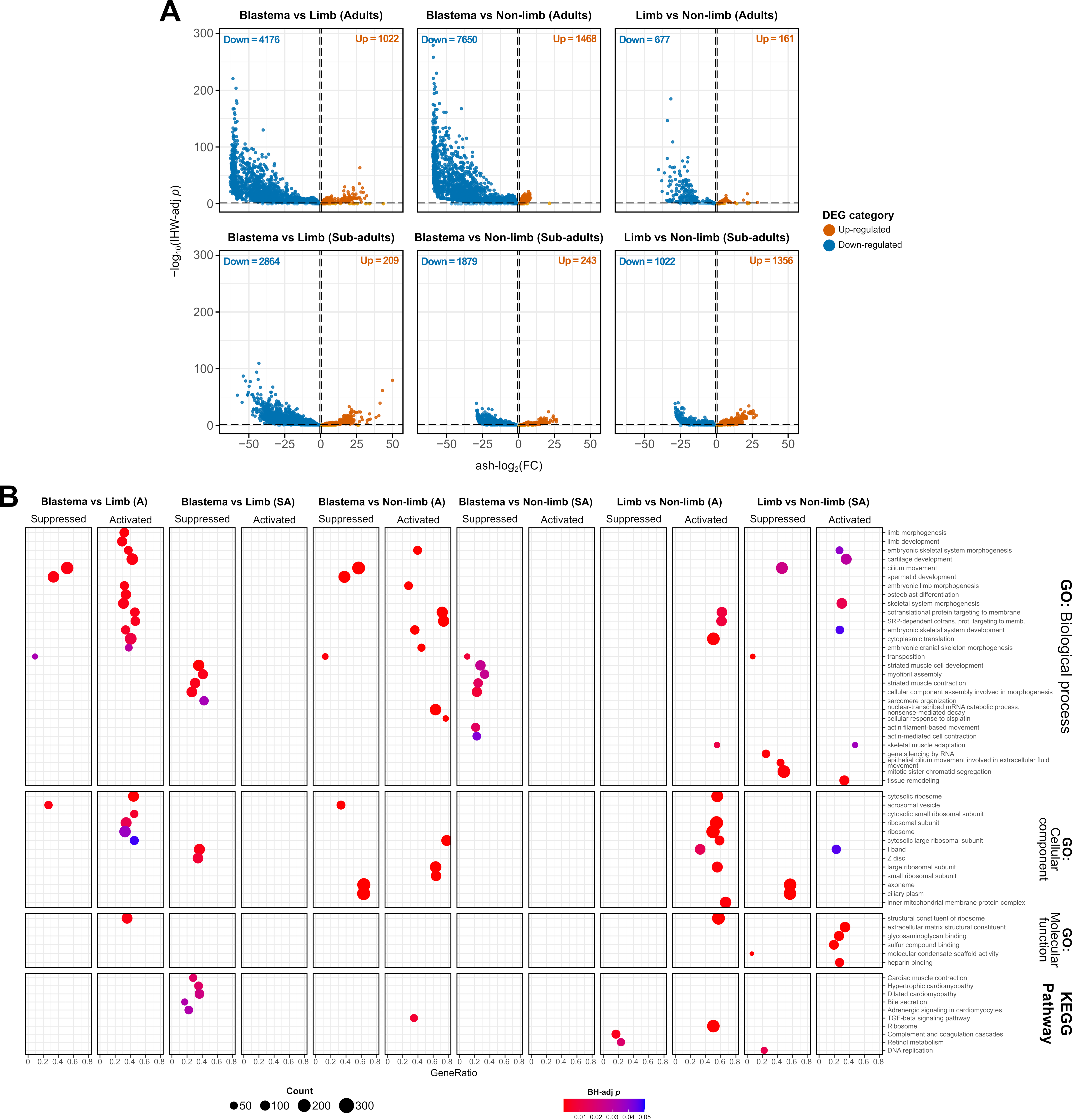

### S3 Fig

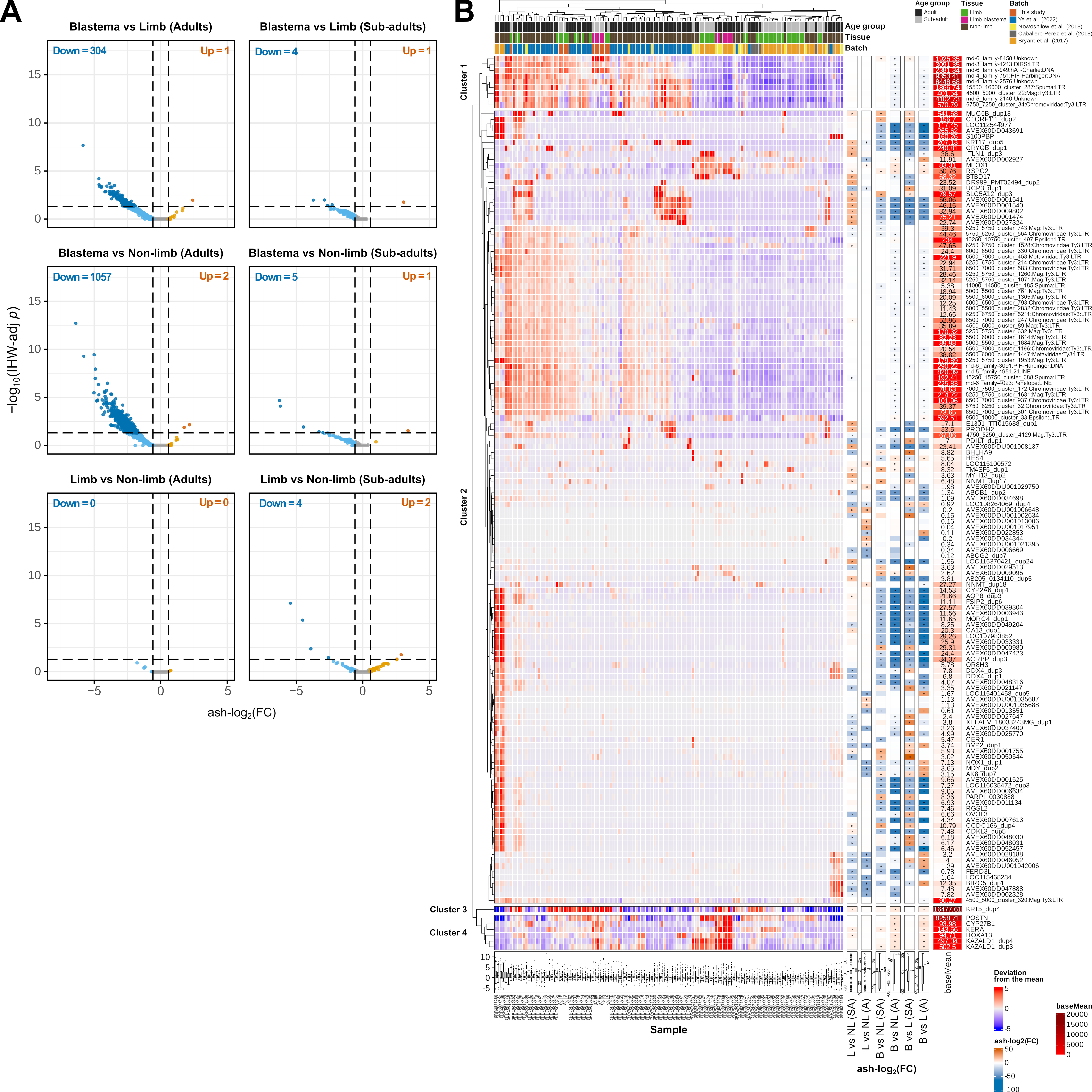
